# A Permutation-Based Framework for Evaluating Bias in Microbiome Differential Abundance Analysis

**DOI:** 10.64898/2026.03.14.711836

**Authors:** Ke Zeng, Anthony Fodor

## Abstract

**Background:** In microbiome research, differential abundance analysis aids in identifying significant differences in microbial taxa across two or more conditions. Statistical approaches used for this purpose include classical tests such as the t-test and Wilcoxon test, as well as methods designed to account for the compositional nature of microbiome data, including ALDEx2, ANCOM-BC2, and metagenomeSeq. In addition, methods originally developed for RNA sequencing data, such as DESeq2 and edgeR, have been frequently applied to microbiome studies. However, the use of these methods has been controversial. One area of concern is whether different modeling frameworks produce accurate p-values when the null hypothesis is true.

**Results:** We evaluated seven methods across six datasets. Four permutation strategies were applied to generate data under the null hypothesis: shuffling sample names, shuffling counts within samples, shuffling counts within taxa, and fully randomizing the counts table. Methods based on the negative binomial distribution (DESeq2 and edgeR) produced p-values that were consistently smaller than expected under the null hypothesis. In contrast, methods that attempt to correct for compositionality (ALDEx2, ANCOM-BC2, and metagenomeSeq) tended to produce larger-than-expected p-values, even when only sample labels were shuffled, a permutation strategy that does not alter compositional structure. These deviations were dependent on dataset characteristics and permutation strategy, suggesting complex interactions between underlying data structure and algorithm performance. Generating data to follow the expected negative binomial distribution did not eliminate the tendency of DESeq2 and edgeR to exaggerate statistical significance. Although similar patterns were observed in RNA sequencing (RNAseq) datasets, the deviations were less pronounced than in microbiome data. In contrast, the classical t-test and Wilcoxon test yielded p-value distributions consistent with theoretical expectations across datasets and permutation strategies.

**Conclusions:** These results indicate that the performance of several widely used differential abundance methods can be problematic under null conditions and may affect biological interpretation. Our findings emphasize the importance of careful method selection and highlight the robustness of simpler statistical approaches for reliable inference.

## BACKGROUND

The study of the microbiome has greatly expanded our understanding of how microbial communities influence environmental processes and host health. A fundamental component of many microbiome investigations is differential abundance analysis (DAA), which seeks to identify microbial taxa whose relative abundances differ between experimental, ecological, or clinical conditions. Despite its central role, DAA remains a statistically challenging problem due to features inherent to microbiome data, including compositionality, sparsity, uneven sequencing depth, and high inter-sample variability. As a result, no single method has emerged as a gold standard that performs optimally across all microbiome study designs, and a wide array of analytical approaches continues to be used in practice.

Classical statistical tests such as the parametric t-test (Student, 1908) and the non-parametric Wilcoxon rank-sum test (Wilcoxon, 1945) have long been applied to microbiome data due to their simplicity and minimal modeling assumptions. In response to the unique characteristics of microbiome count data, several methods have been developed specifically for microbial DAA. These include ALDEx2 (Fernandes et al., 2014), which applies log-ratio transformation combined with Monte Carlo sampling; ANCOM-BC2 (Lin & Peddada, 2024), which corrects for biases introduced by varying sampling fractions; and metagenomeSeq (Paulson et al., 2013), which uses cumulative sum scaling (CSS) normalization to reduce compositional effects. In parallel, methods originally developed for RNAseq data – most notably DESeq2 (Love et al., 2014) and edgeR (Robinson & Smyth, 2007) – have been widely adopted for microbiome studies because they explicitly model count data using the negative binomial distribution and incorporate variance stabilization (Table 1).

**Table 1.**
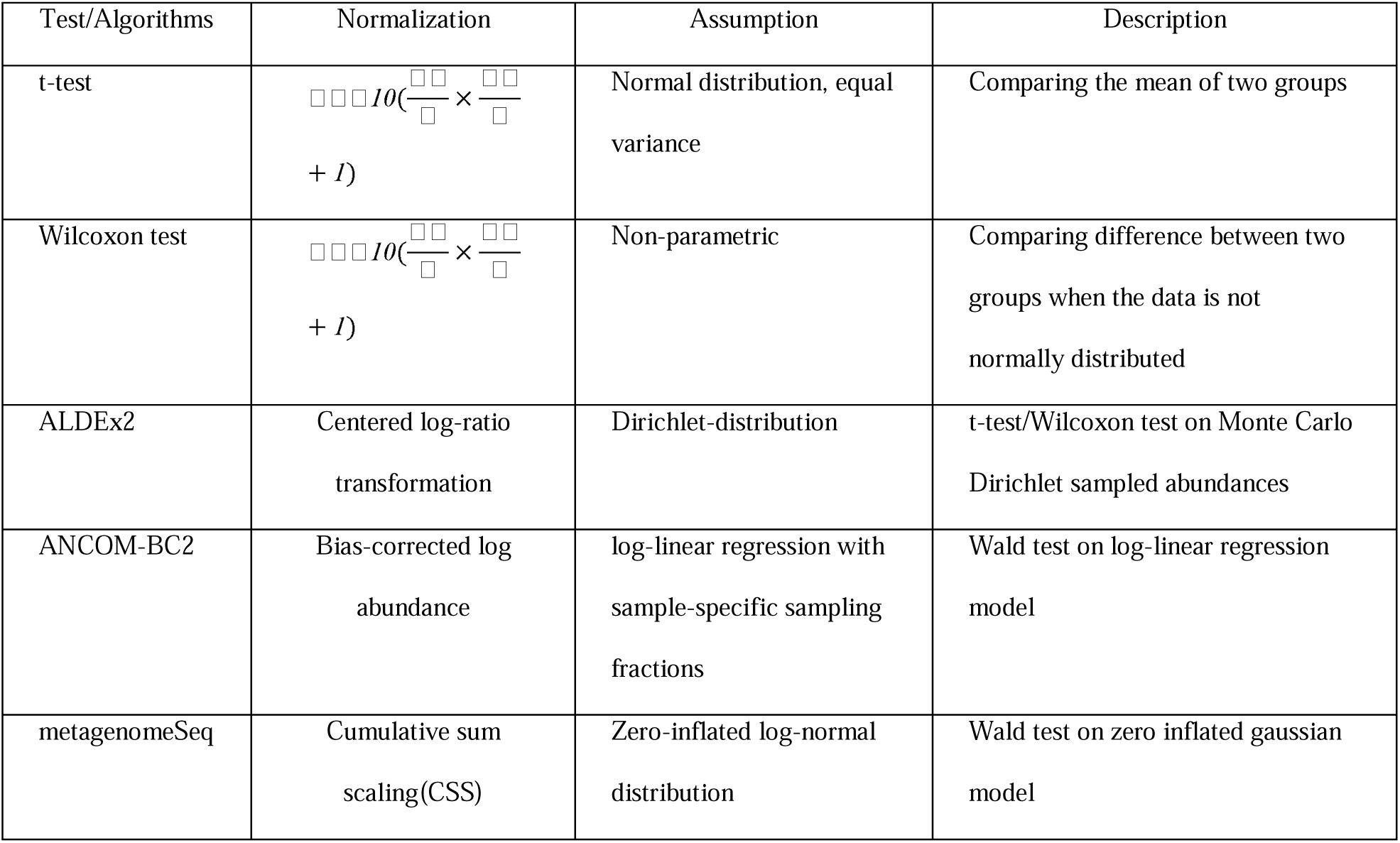
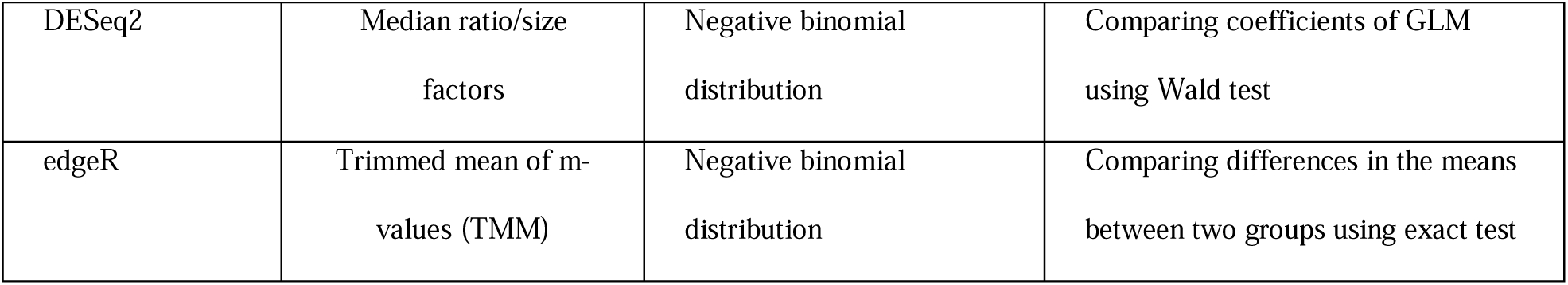
Summary of Differential Abundance Analysis (DAA) Methods. Comparison of the seven methods for DAA: t-test, Wilcoxon test, ALDEx2, ANCOM-BC2, metagenomeSeq, DESeq2, and edgeR,. The methods are compared based on their normalization strategies, assumptions about the underlying data, and their specific approaches for comparing groups.

The appropriateness of RNAseq-derived methods for microbiome DAA has been the subject of ongoing debate. Early work suggested that these approaches could be effectively repurposed for microbiome data (McMurdie & Holmes, 2014), and some comparative evaluations reported that DESeq2 and edgeR performed comparably to microbiome-specific tools, in terms of power and false discovery control (Calgoro et al., 2020). However, more recent studies have raised concerns about their behavior under realistic microbiome null scenarios, reporting poor calibration and elevated false positive rates (Nearing et al., 2022; Hawinkel et al., 2017). Additional evaluations have further suggested that complex parametric methods might yield less reproducible results than simpler, rank-based approaches, with tests such as Wilcoxon rank-sum test often producing more consistent findings than DESeq2 or edgeR (Pelto et al., 2025). Together, these findings show the need for systematic evaluation of DAA methods under well-defined null conditions that reflect realistic properties of microbiome data.

Permutation-based approaches are widely used in microbiome studies to assess statistical significance without relying on strict parametric assumptions. For example, the PERMANOVA assess whether microbial community compositions differ between groups by repeatedly shuffling the group labels assigned to samples and comparing the observed between-group differences to those expected by chance, while other tools like ANCOM (Maldal et al., 2015) and MaAsLin2 (Mallick et al., 2021) have also incorporated permutation schemes for robust inference. Inspired by these practices, in this study, we used four distinct permutation procedures: (i) shuffling the sample labels (e.g., case versus control), (ii) shuffling counts within each sample, (iii) shuffling counts within each taxon, and (iv) shuffling the entire counts table. Each of these permutations produces a dataset in which all the null hypotheses should be true, but each permutation scheme disrupts the underlying structure of the data in different ways. Notably, mislabeling of sample groups – whether due to manmade error or data processing mistakes – can be seen as an unintentional but realistic form of permutation that can influence downstream analyses, motivating the need to understand how these methods respond to such disturbance.

In this study, we evaluate DAA methods across six datasets, including three 16s rRNA microbiome datasets, one whole genome sequencing (WGS) microbiome dataset, and two RNAseq gene expression datasets, to provide a comprehensive comparison. We evaluated the performance of seven commonly used methods: t-test, Wilcoxon test, DESeq2, edgeR, ALDEx2, ANCOM-BC2, and metagenomeSeq. By including taxonomic classification results from both the RDP Classifier and the DADA2 package (Callahan et al., 2016), we further evaluate the extent to which upstream taxonomic classification influences DAA results, and by including different types of datasets, we show that the results can also be dataset – and dataset type – dependent. We found that the t-test and Wilcoxon test are relatively robust to the permutation methods and the type of the dataset, whereas more complex methods frequently fail to produce uniformly distributed p-values under null conditions. ALDEx2 and metagenomeSeq tend to be overly conservative, producing larger p-values under null conditions. ANCOM-BC2, DESeq2, and edgeR tend to exaggerate significance by producing p-values that are too small, potentially leading to false positives for the purpose of DAA.

## METHODS

This study aimed to evaluate the performance of commonly used DA methods and their calibration of p-values under null conditions. We conducted a comparative methodological analysis using six datasets. The study design consisted of applying permutation-based null simulations and distribution-based resampling approaches to assess statistical performance.Three publicly available 16S rRNA datasets at the genus level were analyzed, including two human gut datasets and one soil dataset. In addition, one WGS dataset from mouse gut and two RNAseq datasets (one plant and one mouse gene expression dataset) were also included. All datasets are associated with metadata with two categories, or the two selected categories that are most relevant (Table 2). For each 16S rRNA dataset, two counts tables were generated: one based on taxonomic classification of sequencing reads using the RDP Classifier, and another based on ASV inference followed by taxonomic assignment against the SILVA database (Version 132) using the DADA2 package. Raw counts were prepared with log-normalization for both t-test and Wilcoxon test, where ALDEx2, ANCOM-BC2, metagenomeSeq, DESeq2 and edgeR have used their corresponding built-in normalization methods (Table 1). Count tables were compared using the seven methods. The two sample t-test was performed by using the “t.test” function in base R. The Wilcoxon test was performed using the “wilcox_test” function from the coin package (Version 1.4.1). Analysis with ALDEx2 was performed using default settings with the ALDEx2 package (Version 1.41.0). Analysis with ANCOM-BC2 was performed using the default setting with the ANCOM-BC2 package (Version 2.11.1). Analysis with metagenomeSeq was performed using the default setting with the metagenomeSeq package (Version 1.51.0). Analysis with DESeq2 was performed using the default setting with the DESeq2 package (Version 1.49.3). Analysis with edgeR was performed with default parameters and the “exactTest” function for comparing two groups with the edgeR package (Version 4.7.3). For each taxon in the counts table, the sample-wise mean and variance were calculated using the “mean” and “var” function in base R. Taxa with variance less than or equal to the mean were excluded, as the negative binomial distribution requires the variance to exceed the mean. For the remaining taxa, the parameters of a negative binomial distribution were estimated as: the dispersion parameter 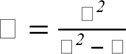, and the success probability 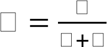, where □ and □^2^ represents the sample mean and variance of the taxon, respectively. New counts values were then generated using the “rnbinom” function in base R, with the number of samples determining the size of the output vector. All analyses were performed using R (Version 4.5.1)

**Table 2.**
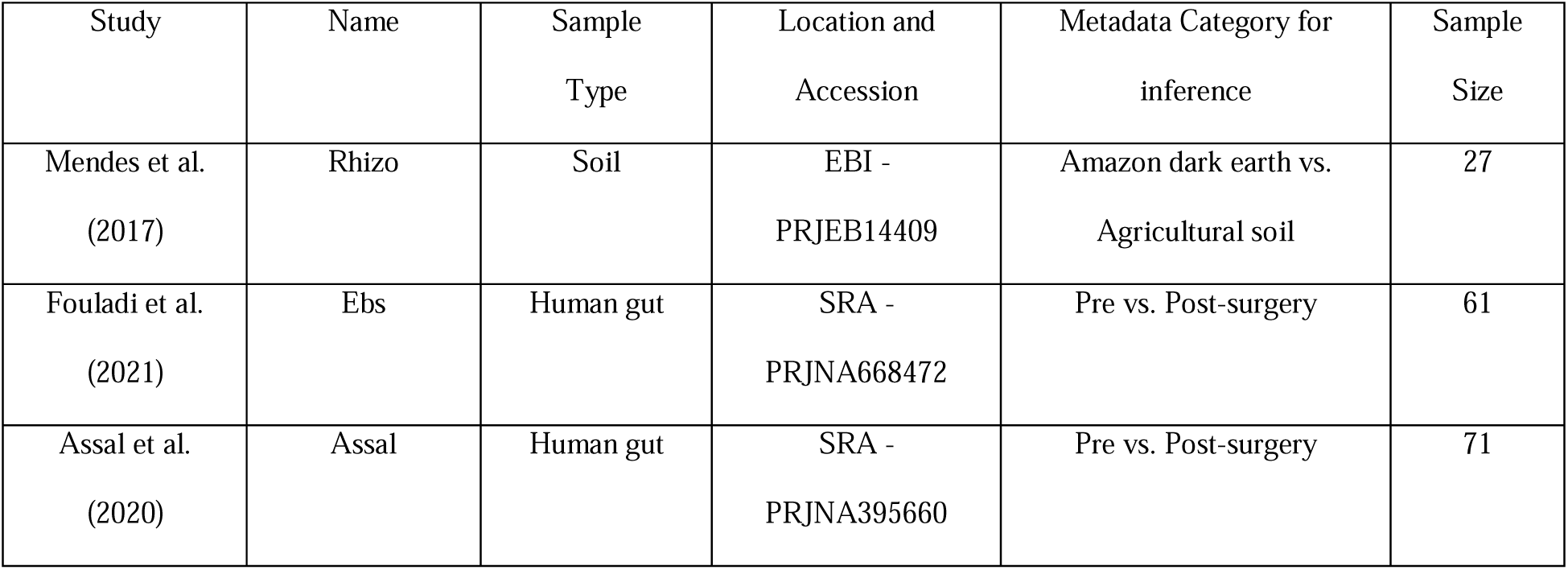
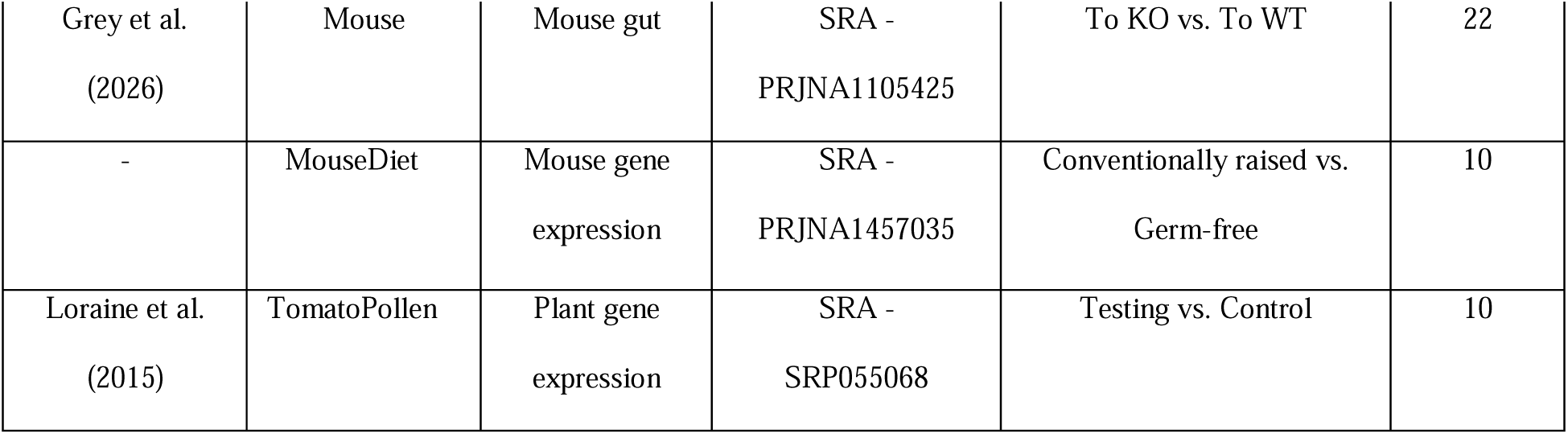
Overview of Datasets Used for Analysis. Summarization of the six datasets used in the analysis, including the study name, sample type, source of sequences, and metadata used for inference.

| Study | Name | Sample Type | Location and Accession | Metadata Category for inference | Sample Size |
| --- | --- | --- | --- | --- | --- |
| Mendes et al.<br>(2017) | Rhizo | Soil | EBI -<br>PRJEB14409 | Amazon dark earth vs.<br>Agricultural soil | 27 |
| Fouladi et al.<br>(2021) | Ebs | Human gut | SRA -<br>PRJNA668472 | Pre vs. Post-surgery | 61 |
| Assal et al.<br>(2020) | Assal | Human gut | SRA -<br>PRJNA395660 | Pre vs. Post-surgery | 71 |
| Grey et al.<br>(2026) | Mouse | Mouse gut | SRA -<br>PRJNA1105425 | To KO vs. To WT | 22 |
| - | MouseDiet | Mouse gene<br>expression | SRA -<br>PRJNA1457035 | Conventionally raised vs.<br>Germ-free | 10 |
| Loraine et al.<br>(2015) | TomatoPollen | Plant gene<br>expression | SRA -<br>SRP055068 | Testing vs. Control | 10 |

## RESULTS

### DESeq2 and edgeR tend to produce smaller p-values compared with t-test and the Wilcoxon test

We performed DAA and differential gene expression analysis, in the case of gene expression counts data, using seven different methods — t-test, Wilcoxon test, DESeq2, edgeR, ALDEx2 (t-test and Wilcoxon test), ANCOMBC2, and metagenomeSeq. These analyses were applied to three different 16S rRNA datasets: two human gut microbiome datasets (assal and bsurgery; Supp. Fig. S1-S2) and one soil dataset (rhizo; Figure 1, Supp. Fig. S3); one WGS mouse gut microbiome dataset (mouse; Supp. Fig. S6), and two RNAseq datasets: one tomato pollen gene expression dataset (tomatoPollen; Supp. Fig. S7) and one mouse gene expression dataset (mouseDiet; Supp. Fig. S8). The log10 p-value for each taxon/gene was adjusted by the direction of the hypothesis for every statistical method, and these results were plotted against each other correspondingly (Figure 1, Supp. Fig. S1-S8).

**Figure 1.**
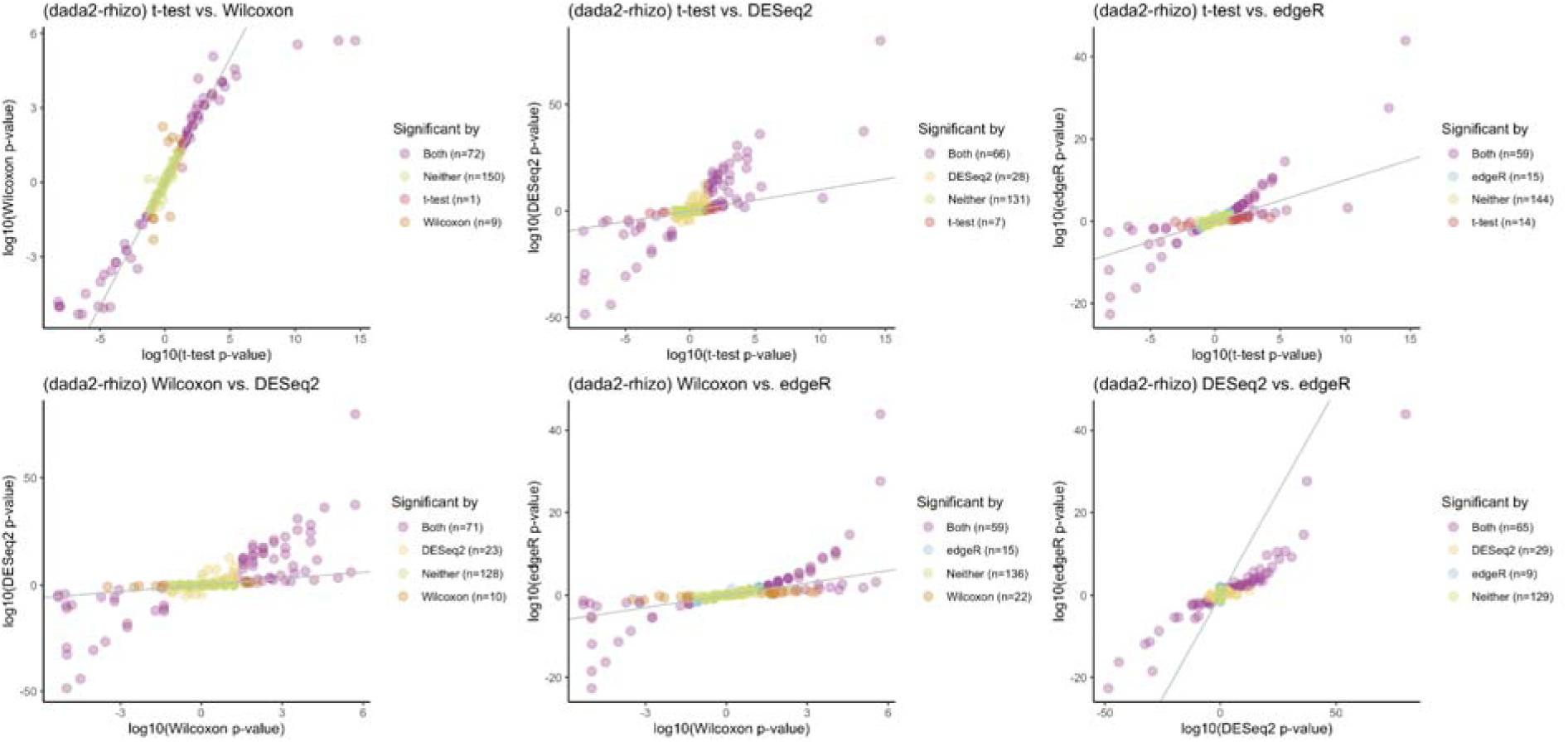
Pairwise comparison of p-values across differential abundance analysis (DAA) methods for the DADA2-rhizo dataset. Each scatter plot compares the log-transformed p-values between two DAA methods, as applied to the same dataset. Each point represents a taxon, and the significant threshold is set at p-value <0.05 (before FDR correction), with colors indicating whether the taxon is identified as significant by both methods, by only one method (Wilcoxon, t-test, DESeq2, or edgeR), or by neither method.

We make the following observations about this analysis:

1. Across all datasets, DESeq2 and edgeR are less conservative for some of the taxa or genes, as evidenced by the production of smaller p-values and higher log10 p-values compared to the t-test, Wilcoxon test, ALDEx2 (t-test and Wilcoxon test), ANCOMBC2, and metagenomeSeq (Supp. Fig. S9-S11).
2. The degree to which DESeq2 and edgeR are less conservative is dataset dependent (Supp. Fig. S1-S3).
3. The degree to which DESeq2 and edgeR are less conservative also varies depending on the pre-processing method used (Figure 1, Supp. Fig. S1-S5).
4. ALDEx2 (t-test and Wilcoxon test) and metagenomeSeq tend to be more conservative than t-test and Wilcoxon test in most datasets (Supp. Fig. S9-S17).
5. ANCOM-BC2 performs similarly to t-test and Wilcoxon test in many cases, but remains more conservative than DESeq2 and edgeR (Supp. Fig. S9-S17).

For the 16S rRNA datasets, we included results from both the RDP classifier, a method that uses a naive Bayesian algorithm to assign sequences to taxonomic groups (Wang et al., 2007), and DADA2, which improves accuracy by correcting sequencing errors and identifying the amplicon sequence variants (ASVs) before classifying them using a reference database (Callahan et al., 2016). Most taxa share similar p-values across methods, but for a few, DESeq2 and edgeR yield p-values that are much smaller, with differences of up to 15-fold in the Rhizo dataset and similar magnitudes in the other datasets (Figure 1, Supp. Fig. S3). ALDEx2 and metagenomeSeq, on the other hand, tend to be much more conservative compared to t-test and Wilcoxon test. ANCOM-BC2 produces less stable results: it is more conservative than DESeq2 and edgeR for most datasets, less conservative than ALDEx2 and metagenomeSeq, and comparable but inconsistent relative to t-test and Wilcoxon test.

We also included the comparison for the WGS dataset and two RNAseq gene expression datasets. These comparisons showed trends consistent with those observed in the 16s rRNA datasets (Supp. Fig. S6-S8, S15-S17). Overall, this trend suggests that DESeq2 and edgeR might be more sensitive when detecting differentially abundant taxa or differentially expressed genes across different types of sequencing data, regardless of the pre-processing method. In contrast, the t-test, Wilcoxon test, ALDEx2, and metagenomeSeq have shown to be more stable and conservative, with p-values clustering closer to the diagonal identity line and in a similar range of value when plotted against each other. ANCOM-BC2 remains less stable, more conservative than DESeq2 and edgeR (if not similar to them), and less conservative than ALDEx2 and metagenomeSeq (Supp. Fig. S15-S17).

### DESeq2 and edgeR yield small p-values that are insensitive to permutations

To test how the correlation and variance structure within the counts table affect the results of DAA from the seven methods, we conducted a series of shuffling experiments. Four different types of permutations were applied to the data:

1. **Sample name tag shuffling**: In this experiment, we shuffled the sample name tags, mixing the labels regardless of the group they belong to, without altering the actual counts data (Figure 2A).
2. **Intra-sample counts shuffling**: We shuffled the counts within each sample, thereby changing the relationship between specific taxa and their respective counts while maintaining the counts data within each sample (Figure 2B).
3. **Intra-taxon counts shuffling**: We shuffled the counts within each taxon, scrambling their distribution across samples while preserving the total counts for each taxon (Figure 2C).
4. **Full counts table shuffling**: The entire counts table was randomized, destroying the original data structure (Figure 2D).

**Figure 2.**
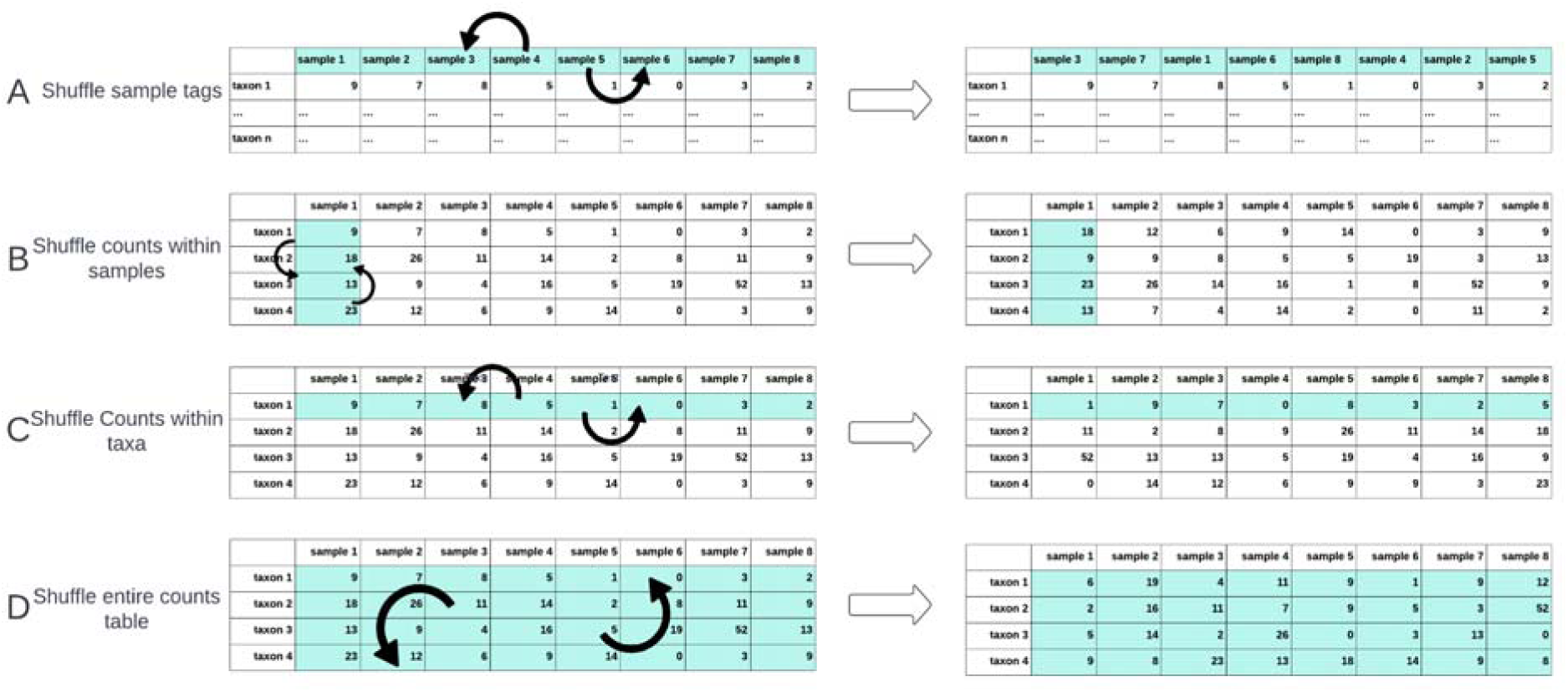
Visualization for four different types of permutation applied to the counts table. Each panel illustrates a different permutation method used to alter the data structure of the counts table before applying differential abundance analysis (DAA).

Each shuffling procedure was repeated 100 times randomly, with DAA performed on each shuffled table using the seven methods. For each shuffled dataset, we calculated the fraction of significant results (p-value < 0.05) produced by each method, and summarized these results using boxplots, in which the y-axis represents the proportion of significant results across 100 shuffles, with a red dashed line at 0.05, indicating the expected fraction under the null hypothesis (Figure 3).

**Figure 3.**
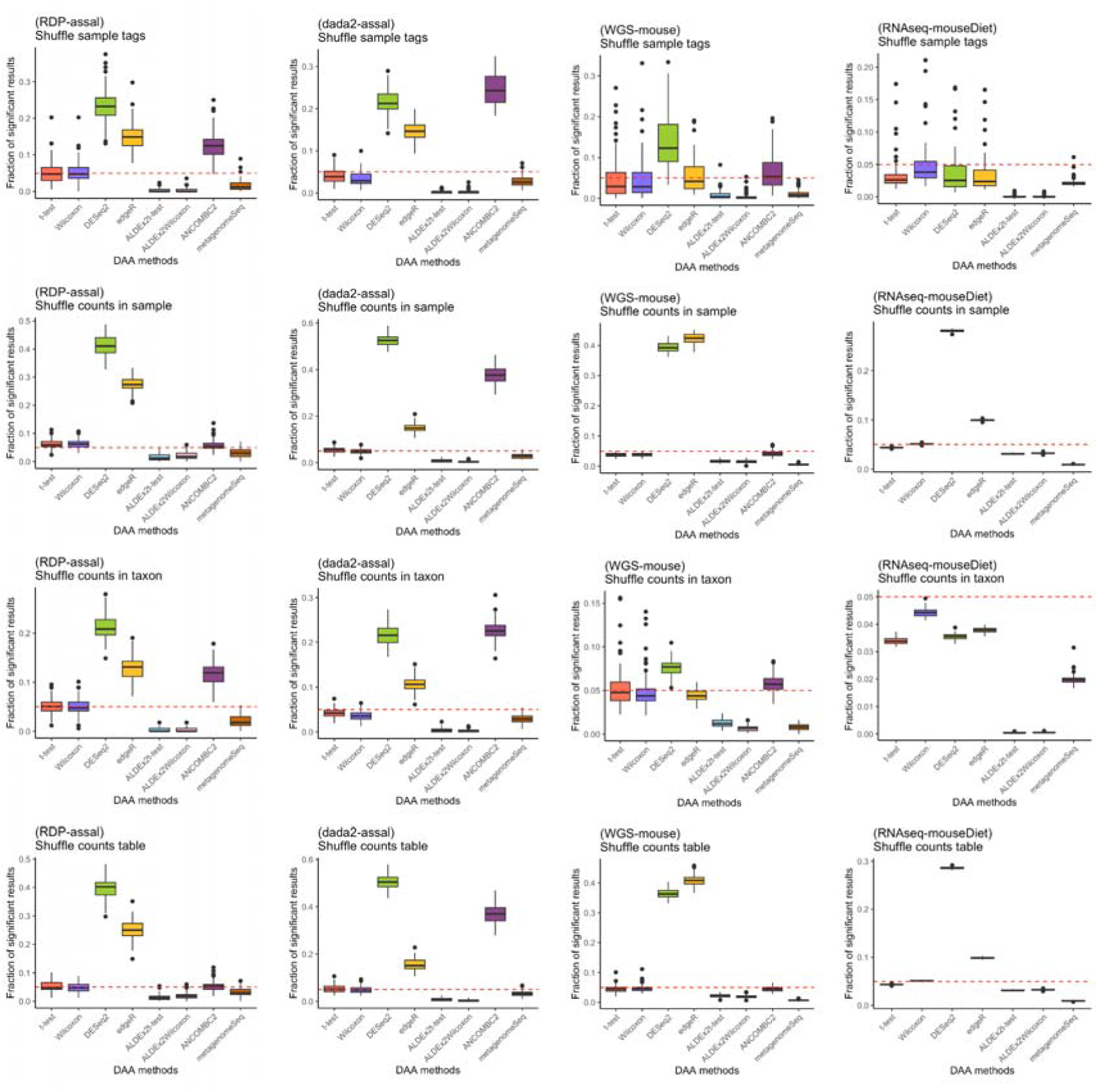
Proportion of significant p-values (p<0.05) generated by seven differential abundance analysis (DAA) methods across four different types of datasets under four shuffling scenarios. Each row shows one shuffling scenario: Shuffle sample tags, shuffle counts in sample, shuffle counts in taxon, and shuffle counts table. Each boxplot shows the distribution of the proportion of significant results from 100 iterations of shuffling from each dataset and DAA method. The dashed red line at 0.05 represents the expected proportion of false positives under a uniform null hypothesis.

We make the following observations about the permutation experiments:

1. Across all the permutation types, the t-test and Wilcoxon test produced fractions of significant results that are close to 0.05 (as expected), aligning with the null hypothesis when group labels or counts are randomized.
2. ALDEx2 (t-test and Wilcoxon test) and MetagenomeSeq are generally conservative with fractions of significant results generally below the 0.05 threshold.
3. DESeq2 often showed higher fractions of significant results than expected, indicating its insensitivity to the permutations.
4. edgeR also showed higher fractions of significant results than expected, but often lower than DESeq2.
5. ANCOM-BC2 shows inconsistent trends of results on different datasets with higher fractions of significant results in some datasets, but not all.
6. The insensitivity to the permutation was dependent on the permutation type and the sequencing/characterization method.

Being insensitive to the permutations indicates that DESeq2, edgeR, and ANCOM-BC2 likely tend to be affected more by the surrounding data structure, and the three methods might detect false positives, even when the true biological signals are removed from the datasets (Figure 3; Supp. Fig. 18). For datasets such as rhizo, when processed by the RDP Classifier, and the two RNAseq datasets, all methods – including DESeq2 and edgeR – produced results close to or below the expected 0.05 threshold for certain shuffling experiments. This shows that some dataset-specific characteristics might have mitigated the methods’ sensitivity to the shuffling experiment (Supp. Fig.18).

### Resampling data from negative binomial distribution can still yield smaller than expected p-values for DESeq2 and edgeR

Because DESeq2 and edgeR assume a negative binomial distribution and showed differing results in the previous experiments, we tested whether deviation from this assumption could explain their tendency to produce the small p-values. To do this, we resampled the counts data across all datasets to explicitly follow a negative binomial distribution, creating a controlled setting that aligns with the assumption with these methods. We then reapplied the four types of shuffling experiments to the resampled data to observe whether this procedure affected the behavior of the methods. For each taxon or gene, the mean and variance of the counts were calculated, and new counts were resampled from a negative binomial distribution using the corresponding mean and variance (Figure 4). By resampling the counts within each taxon, the variance of the counts was equalized, and the overall structure of the dataset was forced into a negative binomial distribution. The shuffling experiments were then performed 100 times randomly for each dataset. For each resampled and shuffled counts table, DAA was performed using the t-test, the Wilcoxon test, DESeq2, and edgeR. The fraction of significant results (p-value < 0.05) was calculated again (Figure 5). The red dashed line at 0.05 indicates the expected threshold under the null hypothesis.

**Figure 4.**
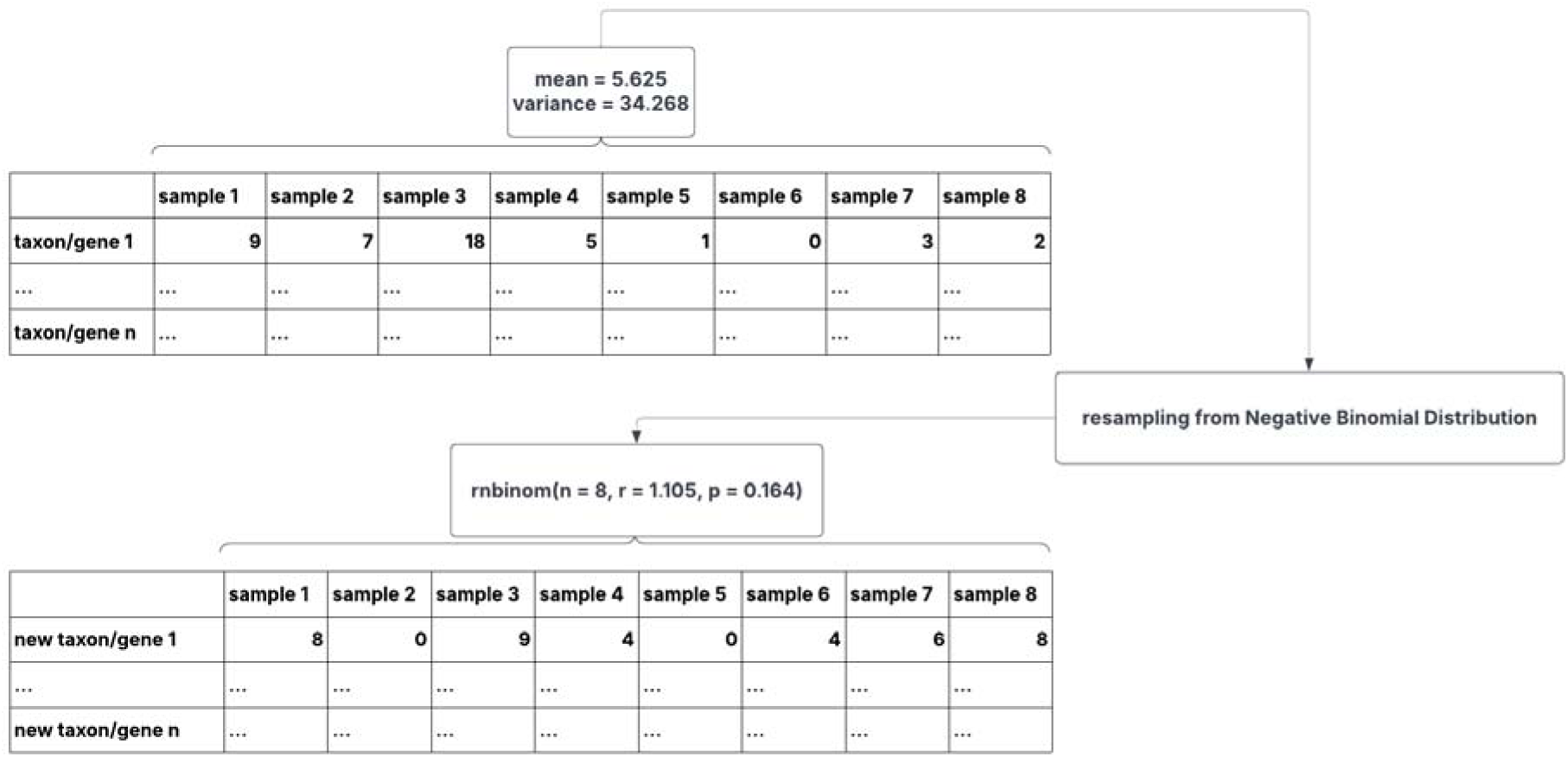
Visualization for the resampling process applied to the counts tables. For each taxon or gene, the mean and variance across samples were calculated. Counts for each taxon or gene were then resampled from this negative binomial distribution, creating a new resampled dataset.

**Figure 5.**
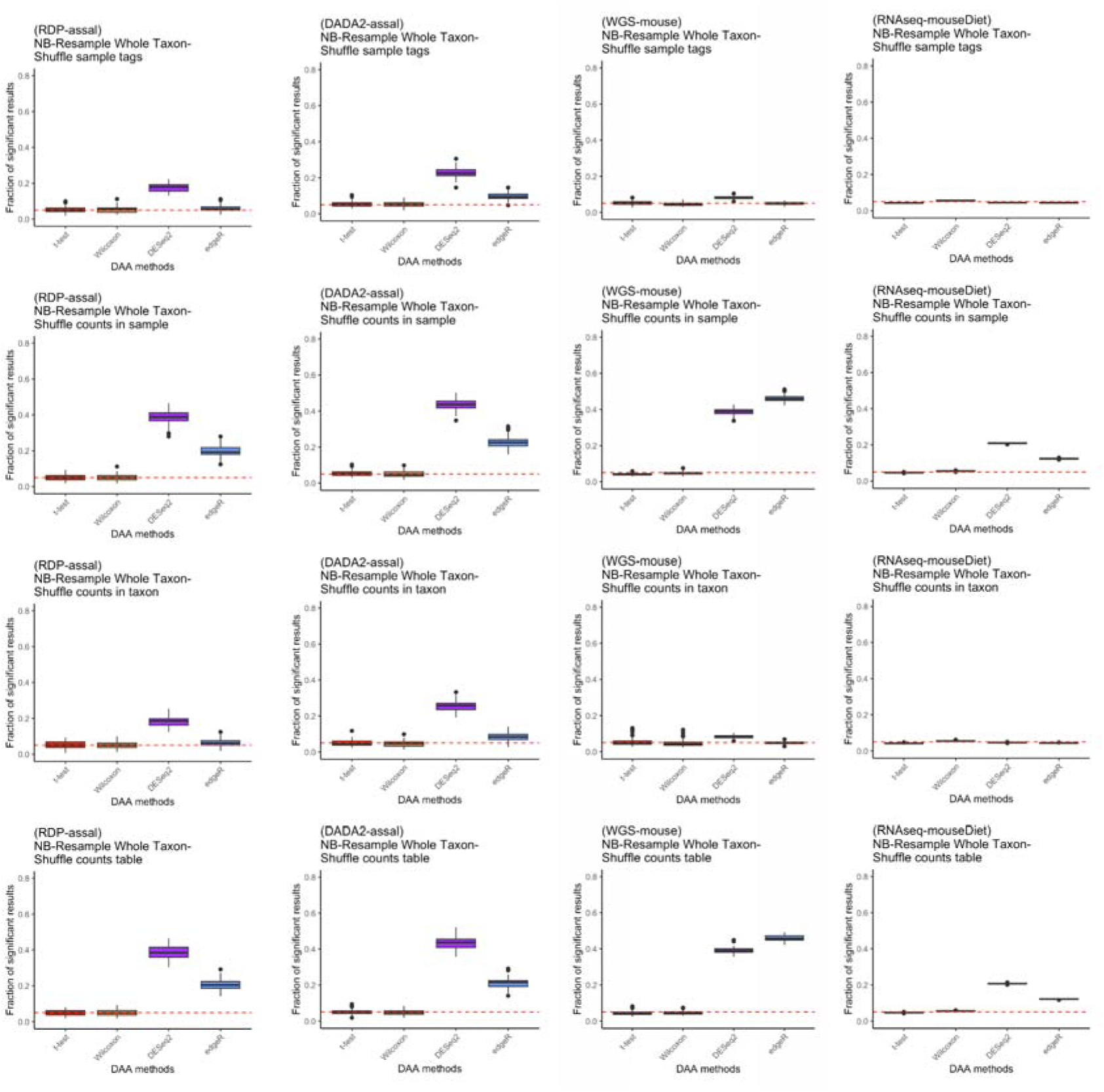
Fraction of significant p-values (p<0.05) from four differential abundance analysis (DAA) methods across four different types of datasets after resampling and shuffling. Resampling was performed 1 time for each datasets, and the resampled datasets were shuffled for 100 times with each shuffling method. Fraction of significant results is shown for each shuffling method. The dashed red line at 0.05 represents the expected proportion of false positives under a uniform null hypothesis.

We make the following observations about this experiment:

1. Both t-test and Wilcoxon test remained robust for the majority of the datasets, as seen previously.
2. In most of the cases, the resampling procedure did not cause DESeq2 and edgeR to become more conservative in the permutation experiments, producing fractions of significant results mostly still above or far from the 0.05 threshold, suggests that the surrounding data structure or some small background signatures might be the factor that affects the results of DESeq2 and edgeR, rather than individual taxa counts not obeying a negative binomial distribution.
3. Occasionally, edgeR produces slightly lower fractions of significant results compared with the non-resampled shuffling experiment results (Figure 5, Supp. Fig. 19).

## DISCUSSION

Across the seven DAA methods evaluated, clear and consistent patterns emerged in their performance under null conditions. The t-test and Wilcoxon test produced p-value distributions that were close to uniform under the null hypothesis, while ALDEx2 and metagenomeSeq tended to be overly conservative, yielding p-values that were systematically larger than expected. While this conservatism could reduce false positives, it also suggests reduced sensitivity and potential loss of power in settings where true biological differences are present. ANCOM-BC2 exhibited intermediate behavior. Although it did not produce the extremely small p-values observed for DESeq2 and edgeR in the log10 p-value experiment, it nevertheless found more significant taxa than expected under several shuffling schemes.

To align the data with the assumptions of DESeq2 and edgeR, we also resampled from the negative binomial distribution. However, our results show that, even under these idealized conditions, these methods continued to yield more than the expected number of significant results. This suggests that their tendency to produce a greater number of significant results is not simply due to lack of fit to parametric assumptions, but rather their reliance on the global variance structure and information shared across the counts table.

We do not fully understand why each algorithm produces unexpected results under null. We speculate that the inflation in ANCOM-BC2 is likely attributable to the inclusion of sparse taxa, which were retained to maintain consistency with the settings used for other methods that did not apply filtering. The presence of sparse features can destabilize standard error estimation in ANCOM-BC2, inflate Wald test statistics, and consequently lead to spurious significance. As for DESeq2 and edgeR, our findings point toward variance stabilization and cross-feature information sharing as the key factors driving the observed artifacts. Although DESeq2 and edgeR are designed to improve statistical power, and sometimes they do produce appropriately uniform p-value distributions under certain shuffled conditions, they are also prone to reporting significant results even when the biological signal has been deliberately removed – something we did not observe among t-test, Wilcoxon test, ALDEx2, or metagenomeSeq results. This raises concerns about the robustness of DESeq2 and edgeR, especially in microbiome datasets where assumptions about count distribution and variance structure might not hold.

The problems we have observed for DESeq2 and edgeR appear to be more pronounced in microbiome data compared to RNAseq datasets. There are several possible reasons: microbiome datasets often contain far fewer features (taxa) than RNAseq datasets have genes, and they tend to exhibit a sparser data structure with a high proportion of zero counts. Moreover, the mean-variance relationship in microbiome data could differ from that in RNAseq, further challenging the assumptions of negative binomial-based models. While we do not know the true cause of these artifacts, our findings suggest that not only microbiome researchers, but also RNAseq users, should apply DESeq2 and edgeR with caution – especially in instances where the data structure deviates from model expectations.

In real world scenarios, such as sample mislabeling or metadata sorting errors – a situation that has occurred in a previous publication, which was since corrected (Kleiman et al., 2019) – methods like DESeq2 and edgeR may still produce convincing, yet misleading, results. Because these methods are not inherently sensitive to such permutations, false positives could potentially go undetected, leading to false biological interpretations. While some of our permutation strategies are artificial and not tied to a specific real world scenario, they still show how these methods can fail to distinguish real signal from random noise.

A recent benchmarking study by Yang and Chen (2022) independently reached a similar conclusion: that DESeq2 and edgeR are prone to inflated false positives in microbiome DAA, especially under compositional and confounded settings. While their study provides broad benchmarking across many methods and scenarios, our work offers a more focused exploration of why these problems occur. Specifically, we extended their findings by applying multiple types of shuffling strategies and demonstrate that the inconsistencies are not due to parametric misfits of the negative binomial model itself, but rather to how these methods borrow variance information across features, which can be problematic in datasets with strong inter-taxon dependencies. An earlier paper (Hawinkel et al., 2017) used a variety of techniques including parametric and non-parametric-based simulations and sub-sampling to argue that DA methods “break the promise” of FDR control. Our results are broadly consistent with their conclusions, but we make the important observation that FDR control can be reliably obtained across a wide range of permutation tests by utilizing the t-test and Wilcoxon test rather than more recently derived methods.

Interestingly, despite employing an orthogonal experiment design focused on replicability across subsampled independent real world datasets rather than controlled permutations, another recent study by Pelto et al. (2025) arrives at a similar conclusion: simpler, non-parametric methods, such as the t-test and the Wilcoxon test, tend to produce more stable and trustworthy results in microbiome DAA. This independent convergence, from two distinct methodological perspectives, further reinforces the concern that more complex models, such as DESeq2 and edgeR, can be overly sensitive to data structure and prone to generating non-reproducible significant findings, particularly for microbiome data. Taken together, these studies make a compelling case that results from the t-test and the Wilcoxon test are stable when looking at either data subset. This is supported in both the Pelto et al. study and in our study when evaluating shuffled datasets in which the null hypothesis is always true.

Although microbiome data are inherently compositional, our results show that compositionality alone does not explain the divergent behaviors observed among the different methods. In some of the permutation strategies used here, the compositional structure, sparsity, and library size distributions were either partially or fully preserved, yet both t-test and Wilcoxon test consistently produced the expected p-value distributions under the null hypothesis. In contrast, DESeq2 and edgeR frequently exhibited right-skewed p-value distributions. Our findings suggest that the main source of bias in DESeq2 and edgeR did not arise from compositionality itself, but from their reliance on global dispersion estimation and cross-taxon information sharing.

We demonstrated substantial study-to-study variation in the performance of many of these algorithms. If researchers wish to use algorithms other than the t-test and Wilcoxon test, one potential application of our study is that researchers could use shuffling-based diagnostic strategies, such as those presented here, to assess the validity of their DAA results. Under such a scheme, researchers could demonstrate that significant results are sensitive to permutation schemes, increasing confidence that assumptions of more complex statistical models did not produce the observed significant results. However, our results suggest that if simpler methods like the t-test and Wilcoxon test can achieve reasonable power, they should be favored due to their interpretability, robustness, and resistance to spurious results under data perturbations.

This study has a few potential limitations. First, the analyses were conducted primarily using the default parameters and settings to reflect common practice in applied research. Although alternative tuning processes could potentially influence the performance of different methods, the persistence of deviations under multiple permutation and resampling procedures suggests that the observed patterns are driven by fundamental modeling characteristics rather than isolated implementation choices. Second, while we examined multiple microbiome and RNAseq datasets, they do not encompass the full spectrum of data sparsity, sequencing depth, or experimental heterogeneity encountered in practice. Third, our experiments were restricted to two group comparisons, and we did not assess performance in multi-factor designs, longitudinal settings, or models incorporating covariates. Method behavior may differ under these more complex scenarios. Nonetheless, the consistency of the findings across datasets, shuffling strategies, and distribution-based resampling indicates that the main conclusions are unlikely to be artifacts of any single dataset or analytical choice.

## CONCLUSION

Across multiple datasets and permutation-based null conditions, negative binomial–based methods frequently produced more significant results than expected, while classical parametric and non-parametric tests yielded p-value distributions consistent with theoretical expectations. Methods designed to account for compositionality exhibited conservative behavior, with varying degrees of deviation depending on dataset and permutation strategy.

This study provides a unified permutation-based framework for assessing differential abundance methods under multiple null scenarios and demonstrates that increased methodological complexity does not necessarily lead to more reliable inference. By directly comparing microbiome-specific approaches, RNAseq-derived methods, and classical statistical tests across multiple dataset types, our findings provide practical guidance for selecting DAA methods that yield robust, interpretable, and reproducible results in microbiome research.

## LIST OF ABBREVIATIONS

DAA: Differential abundance analysis
TMM: Trimmed mean of M-values
OTU: Operational taxonomic unit
FDR: False discovery rate
DA: Differential abundance
PERMANOVA: Permutational multivariate analysis of variance
RPKM: Reads per kilobase of transcript per million reads mapped
rRNA: Ribosomal RNA
GLM: Generalized linear model
WGS: Whole genome sequencing
RNAseq: RNA sequencing
CSS: Cumulative sum scaling
ASVs: Amplicon sequence variants

## DECLARATIONS

### Ethics approval and consent to participate

Not applicable.

### Consent for publication

Not applicable.

### Availability of data and materials

The datasets used in this study are all publicly available. The availability and source of raw sequences for each dataset are summarized in Table 2. R scripts used in this study are available at GitHub (https://github.com/kezengke/DAalgorithms). Additional requests and questions can be addressed to KZ.

### Authors’ contributions

KZ performed the data analysis and drafted the manuscript. KZ and AF contributed to manuscript writing and revisions. AF assisted with testing and provided supervision. All authors read and approved the final manuscript.

## Acknowledgements

The authors thank the members of the Fodor lab for all the help and discussions.

AI tools were used to improve the readability and clarity of the manuscript. In addition, AI assistance was used to generate testing code for the RDP-Rhizo dataset in both R and Python. The AI-generated scripts produced results consistent with those obtained from the original manually written code (Supplementary Fig. S20).

## REFERENCES

Student. (1908). The Probable Error of a Mean. Biometrika, 6(1), 1–25. 10.2307/2331554

Wilcoxon, F. (1945). Individual Comparisons by Ranking Methods. Biometrics Bulletin, 1(6), 80–83. 10.2307/3001968

Fernandes, A. D., Reid, J. N., Macklaim, J. M., McMurrough, T. A., Edgell, D. R., & Gloor, G. B. (2014). Unifying the analysis of high-throughput sequencing datasets: characterizing RNA-seq, 16S rRNA gene sequencing and selective growth experiments by compositional data analysis. Microbiome, 2, 15. 10.1186/2049-2618-2-15

Lin, H., Peddada, S.D. Multigroup analysis of compositions of microbiomes with covariate adjustments and repeated measures. Nat Methods 21, 83–91 (2024). 10.1038/s41592-023-02092-7

Paulson, J., Stine, O., Bravo, H. et al. Differential abundance analysis for microbial marker-gene surveys. Nat Methods 10, 1200–1202 (2013). 10.1038/nmeth.2658

Love, M. I., Huber, W., & Anders, S. (2014). Moderated estimation of fold change and dispersion for RNA-seq data with DESeq2. Genome Biology, 15(12). 10.1186/s13059-014-0550-8

Robinson, M. D., & Smyth, G. K. (2007). Moderated statistical tests for assessing differences in tag abundance. Bioinformatics, 23(21), 2881–2887. 10.1093/bioinformatics/btm453

McMurdie, P. J., & Holmes, S. (2014). Waste Not, Want Not: Why Rarefying Microbiome Data Is Inadmissible. PLOS Computational Biology, 10(4), e1003531. 10.1371/journal.pcbi.1003531

Calgaro, M., Romualdi, C., Waldron, L. et al. Assessment of statistical methods from single cell, bulk RNA-seq, and metagenomics applied to microbiome data. Genome Biol 21, 191 (2020). 10.1186/s13059-020-02104-1

Nearing, J. T., Douglas, G. M., Hayes, M. G., MacDonald, J., Desai, D. K., Allward, N., Casey, Wright, R. J., Dhanani, A. S., Comeau, A. M., & Morgan. (2022). Microbiome differential abundance methods produce different results across 38 datasets. Nature Communications, 13(1), 1–16. 10.1038/s41467-022-28034-z

Stijn Hawinkel, Federico Mattiello, Luc Bijnens, Olivier Thas, A broken promise: microbiome differential abundance methods do not control the false discovery rate, Briefings in Bioinformatics, Volume 20, Issue 1, January 2019, Pages 210–221, 10.1093/bib/bbx104

Pelto, J., Auranen, K., Kujala, J. V., & Lahti, L. (2025). Elementary methods provide more replicable results in microbial differential abundance analysis. Briefings in Bioinformatics, 26(2). 10.1093/bib/bbaf130

Mandal, S., Van Treuren, W., White, R. A., Eggesbø, M., Knight, R., & Peddada, S. D. (2015). Analysis of composition of microbiomes: a novel method for studying microbial composition. Microbial ecology in health and disease, 26, 27663. 10.3402/mehd.v26.27663

Mallick, H., Rahnavard, A., McIver, L. J., Ma, S., Zhang, Y., Nguyen, L. H., … Huttenhower, C. (2021). Multivariable association discovery in population-scale meta-omics studies. PLoS Computational Biology, 17(11), e1009442–e1009442. 10.1371/journal.pcbi.1009442

Callahan, B. J., McMurdie, P. J., Rosen, M. J., Han, A. W., Johnson, A. J. A., & Holmes, S. P. (2016). DADA2: High-resolution sample inference from Illumina amplicon data. Nature Methods, 13(7), 581–583. 10.1038/nmeth.3869

Mendes, L. W., Raaijmakers, J. M., de Hollander, M., Mendes, R., & Tsai, S. M. (2017). Influence of resistance breeding in common bean on rhizosphere microbiome composition and function. The ISME Journal, 12(1), 212–224. 10.1038/ismej.2017.158

Fouladi, F., Carroll, I. M., Sharpton, T. J., Bulik-Sullivan, E., Heinberg, L., Steffen, K. J., & Fodor, A. A. (2021). A microbial signature following bariatric surgery is robustly consistent across multiple cohorts. Gut Microbes, 13(1). 10.1080/19490976.2021.1930872

Wang, Q., Garrity, G. M., Tiedje, J. M., & Cole, J. R. (2007). Naive Bayesian classifier for rapid assignment of rRNA sequences into the new bacterial taxonomy. Applied and environmental microbiology, 73(16), 5261–5267. 10.1128/AEM.00062-07

Callahan, B., McMurdie, P., Rosen, M. et al. DADA2: High-resolution sample inference from Illumina amplicon data. Nat Methods 13, 581–583 (2016). 10.1038/nmeth.3869

Kleiman, S. C., Bulik-Sullivan, E. C., Glenny, E. M., Zerwas, S. C., Huh, E. Y., Tsilimigras, M. C. B., Fodor, A. A., Bulik, C. M., & Carroll, I. M. (2019). Correction: The Gut-Brain Axis in Healthy Females: Lack of Significant Association between Microbial Composition and Diversity with Psychiatric Measures. PLOS ONE, 14(8), e0221724. 10.1371/journal.pone.0221724

Yang, L., & Chen, J. (2023). Benchmarking differential abundance analysis methods for correlated microbiome sequencing data. Briefings in Bioinformatics, 24(1). 10.1093/bib/bbac607

