## supplementary figures for "A Permutation-Based Framework for Evaluating Bias in Microbiome Differential Abundance Analysis"

S1.

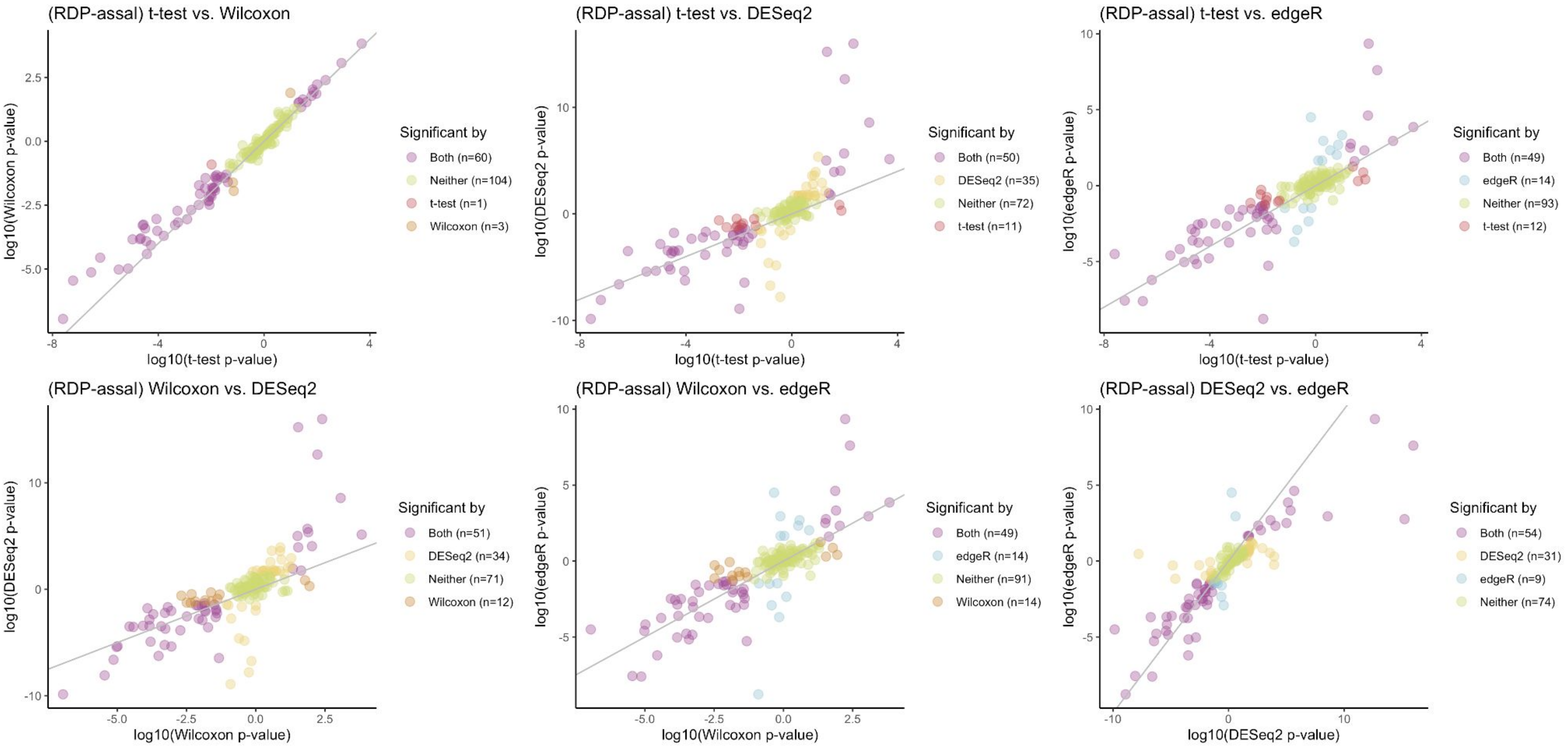
(RDP-assal) t-test vs. Wilcoxon
log10(Wilcoxon p-value)
log10(t-test p-value)
Significant by
Both (n=60)
Neither (n=104)
t-test (n=1)
Wilcoxon (n=3)
(RDP-assal) t-test vs. DESeq2
log10(DESeq2 p-value)
log10(t-test p-value)
Significant by
Both (n=50)
DESeq2 (n=35)
Neither (n=72)
t-test (n=11)
(RDP-assal) t-test vs. edgeR
log10(edgeR p-value)
log10(t-test p-value)
Significant by
Both (n=49)
edgeR (n=14)
Neither (n=93)
t-test (n=12)
(RDP-assal) Wilcoxon vs. DESeq2
log10(DESeq2 p-value)
log10(Wilcoxon p-value)
Significant by
Both (n=51)
DESeq2 (n=34)
Neither (n=71)
Wilcoxon (n=12)
(RDP-assal) Wilcoxon vs. edgeR
log10(edgeR p-value)
log10(Wilcoxon p-value)
Significant by
Both (n=49)
edgeR (n=14)
Neither (n=91)
Wilcoxon (n=14)
(RDP-assal) DESeq2 vs. edgeR
log10(edgeR p-value)
log10(DESeq2 p-value)
Significant by
Both (n=54)
DESeq2 (n=31)
edgeR (n=9)
Neither (n=74)

S2.

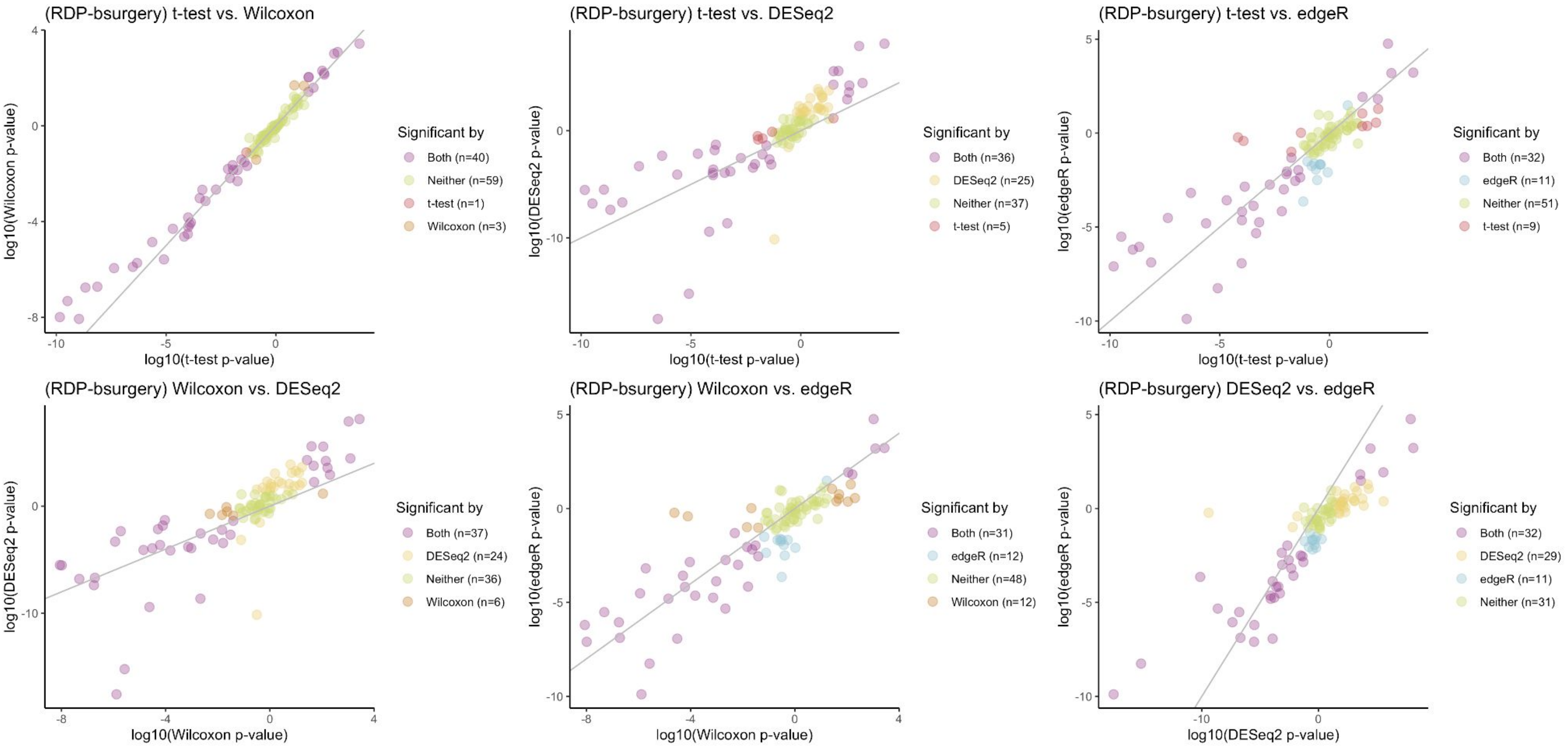
(RDP-bsurgery) t-test vs. Wilcoxon
log10(Wilcoxon p-value)
log10(t-test p-value)
Significant by
Both (n=40)
Neither (n=59)
t-test (n=1)
Wilcoxon (n=3)
(RDP-bsurgery) t-test vs. DESeq2
log10(DESeq2 p-value)
log10(t-test p-value)
Significant by
Both (n=36)
DESeq2 (n=25)
Neither (n=37)
t-test (n=5)
(RDP-bsurgery) t-test vs. edgeR
log10(edgeR p-value)
log10(t-test p-value)
Significant by
Both (n=32)
edgeR (n=11)
Neither (n=51)
t-test (n=9)
(RDP-bsurgery) Wilcoxon vs. DESeq2
log10(DESeq2 p-value)
log10(Wilcoxon p-value)
Significant by
Both (n=37)
DESeq2 (n=24)
Neither (n=36)
Wilcoxon (n=6)
(RDP-bsurgery) Wilcoxon vs. edgeR
log10(edgeR p-value)
log10(Wilcoxon p-value)
Significant by
Both (n=31)
edgeR (n=12)
Neither (n=48)
Wilcoxon (n=12)
(RDP-bsurgery) DESeq2 vs. edgeR
log10(edgeR p-value)
log10(DESeq2 p-value)
Significant by
Both (n=32)
DESeq2 (n=29)
edgeR (n=11)
Neither (n=31)

S3.

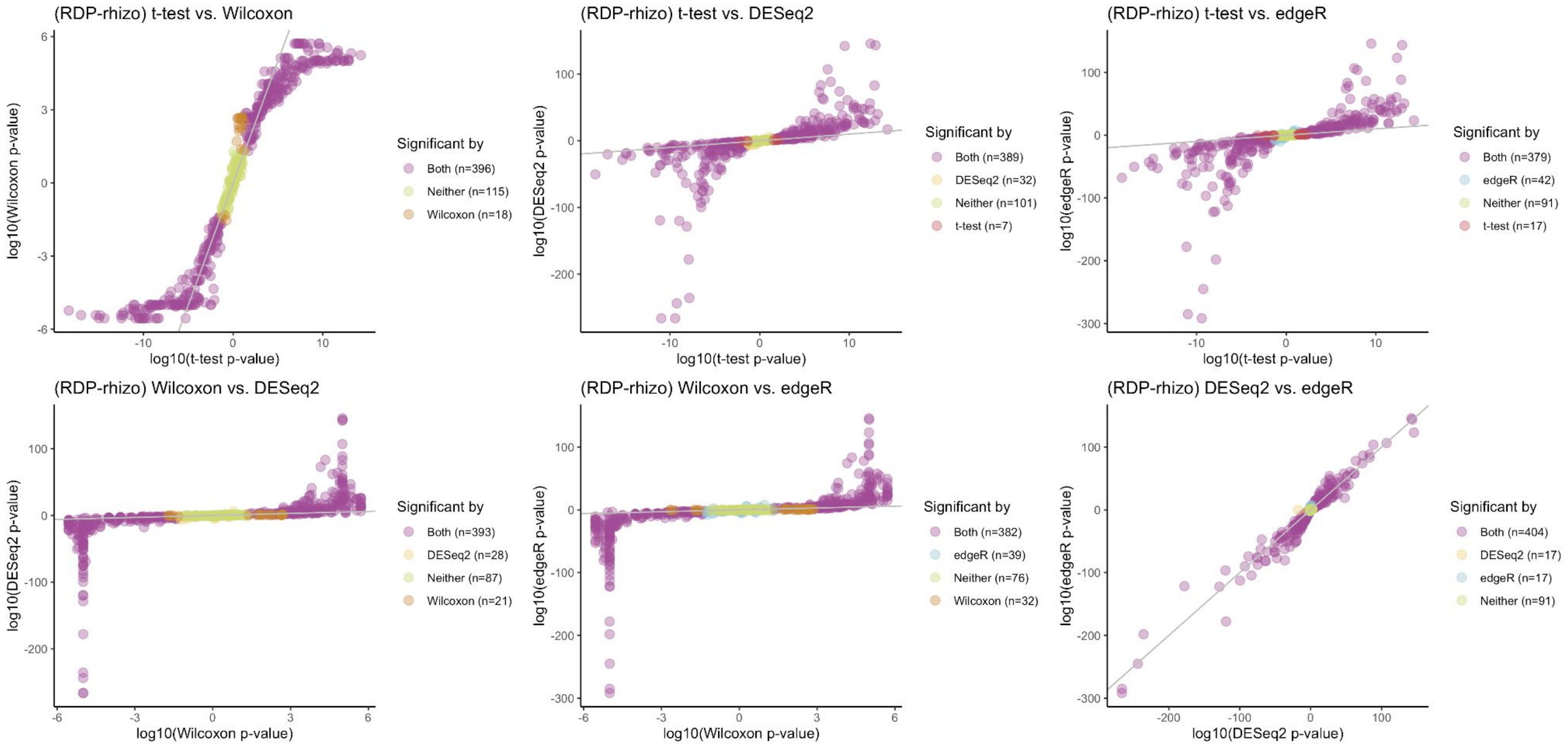
(RDP-rhizo) t-test vs. Wilcoxon
log10(Wilcoxon p-value)
log10(t-test p-value)
Significant by
Both (n=396)
Neither (n=115)
Wilcoxon (n=18)
(RDP-rhizo) t-test vs. DESeq2
log10(DESeq2 p-value)
log10(t-test p-value)
Significant by
Both (n=389)
DESeq2 (n=32)
Neither (n=101)
t-test (n=7)
(RDP-rhizo) t-test vs. edgeR
log10(edgeR p-value)
log10(t-test p-value)
Significant by
Both (n=379)
edgeR (n=42)
Neither (n=91)
t-test (n=17)
(RDP-rhizo) Wilcoxon vs. DESeq2
log10(DESeq2 p-value)
log10(Wilcoxon p-value)
Significant by
Both (n=393)
DESeq2 (n=28)
Neither (n=87)
Wilcoxon (n=21)
(RDP-rhizo) Wilcoxon vs. edgeR
log10(edgeR p-value)
log10(Wilcoxon p-value)
Significant by
Both (n=382)
edgeR (n=39)
Neither (n=76)
Wilcoxon (n=32)
(RDP-rhizo) DESeq2 vs. edgeR
log10(edgeR p-value)
log10(DESeq2 p-value)
Significant by
Both (n=404)
DESeq2 (n=17)
edgeR (n=17)
Neither (n=91)

S4.

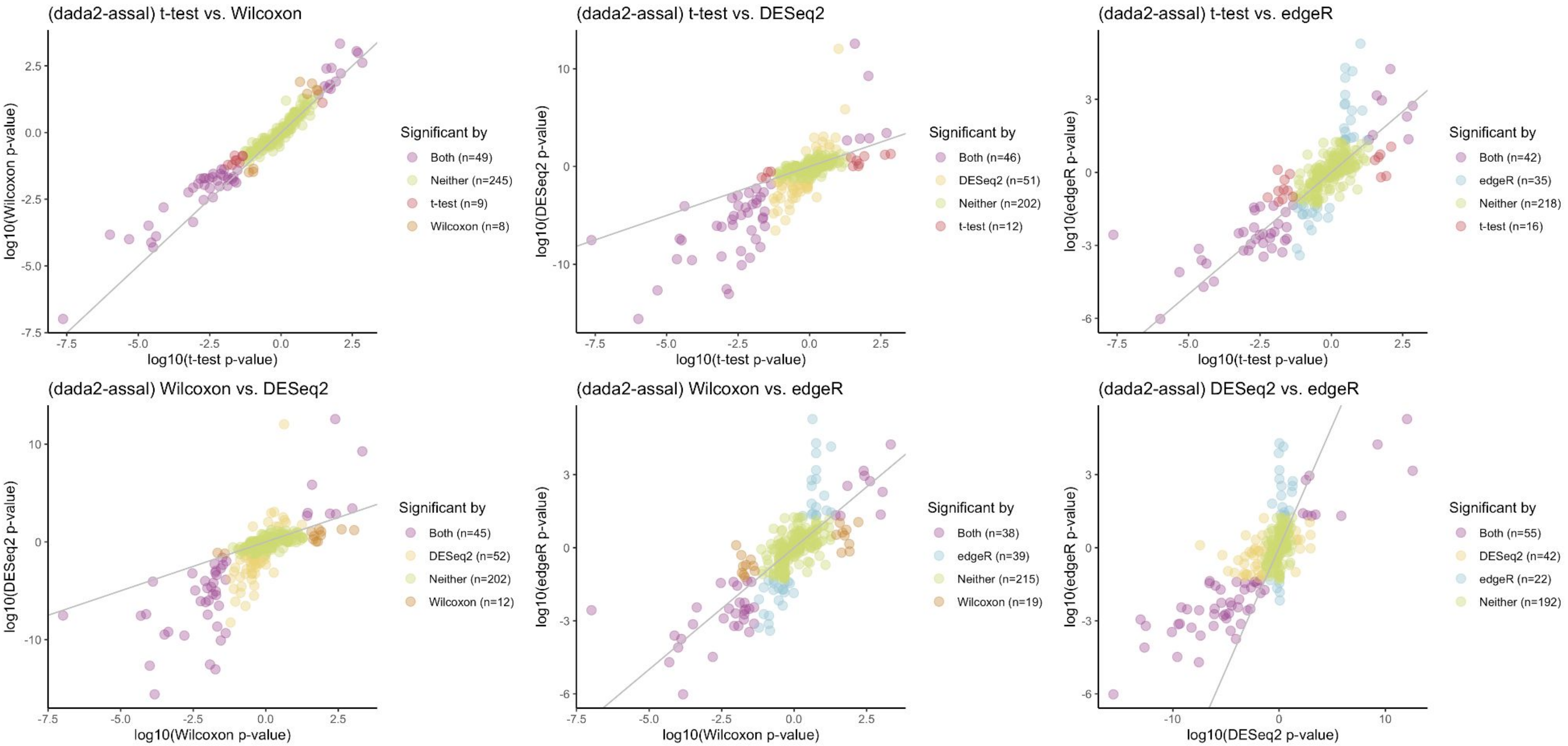
(dada2-assal) t-test vs. Wilcoxon
log10(Wilcoxon p-value)
log10(t-test p-value)
Significant by
Both (n=49)
Neither (n=245)
t-test (n=9)
Wilcoxon (n=8)
(dada2-assal) t-test vs. DESeq2
log10(DESeq2 p-value)
log10(t-test p-value)
Significant by
Both (n=46)
DESeq2 (n=51)
Neither (n=202)
t-test (n=12)
(dada2-assal) t-test vs. edgeR
log10(edgeR p-value)
log10(t-test p-value)
Significant by
Both (n=42)
edgeR (n=35)
Neither (n=218)
t-test (n=16)
(dada2-assal) Wilcoxon vs. DESeq2
log10(DESeq2 p-value)
log10(Wilcoxon p-value)
Significant by
Both (n=45)
DESeq2 (n=52)
Neither (n=202)
Wilcoxon (n=12)
(dada2-assal) Wilcoxon vs. edgeR
log10(edgeR p-value)
log10(Wilcoxon p-value)
Significant by
Both (n=38)
edgeR (n=39)
Neither (n=215)
Wilcoxon (n=19)
(dada2-assal) DESeq2 vs. edgeR
log10(edgeR p-value)
log10(DESeq2 p-value)
Significant by
Both (n=55)
DESeq2 (n=42)
edgeR (n=22)
Neither (n=192)

S5.

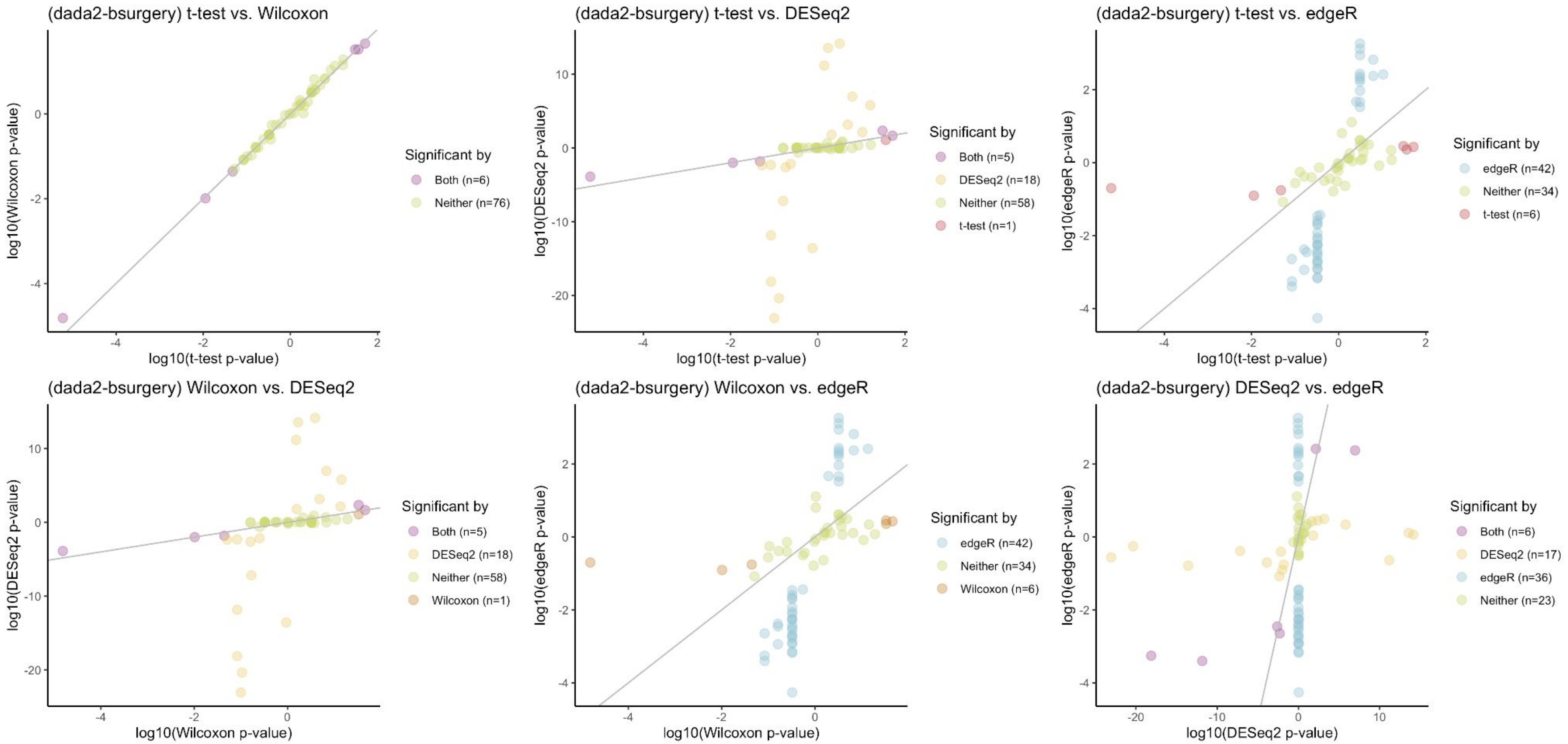
(dada2-bsurgery) t-test vs. Wilcoxon
log10(Wilcoxon p-value)
log10(t-test p-value)
Significant by
Both (n=6)
Neither (n=76)
(dada2-bsurgery) t-test vs. DESeq2
log10(DESeq2 p-value)
log10(t-test p-value)
Significant by
Both (n=5)
DESeq2 (n=18)
Neither (n=58)
t-test (n=1)
(dada2-bsurgery) t-test vs. edgeR
log10(edgeR p-value)
log10(t-test p-value)
Significant by
edgeR (n=42)
Neither (n=34)
t-test (n=6)
(dada2-bsurgery) Wilcoxon vs. DESeq2
log10(DESeq2 p-value)
log10(Wilcoxon p-value)
Significant by
Both (n=5)
DESeq2 (n=18)
Neither (n=58)
Wilcoxon (n=1)
(dada2-bsurgery) Wilcoxon vs. edgeR
log10(edgeR p-value)
log10(Wilcoxon p-value)
Significant by
edgeR (n=42)
Neither (n=34)
Wilcoxon (n=6)
(dada2-bsurgery) DESeq2 vs. edgeR
log10(edgeR p-value)
log10(DESeq2 p-value)
Significant by
Both (n=6)
DESeq2 (n=17)
edgeR (n=36)
Neither (n=23)

S6.

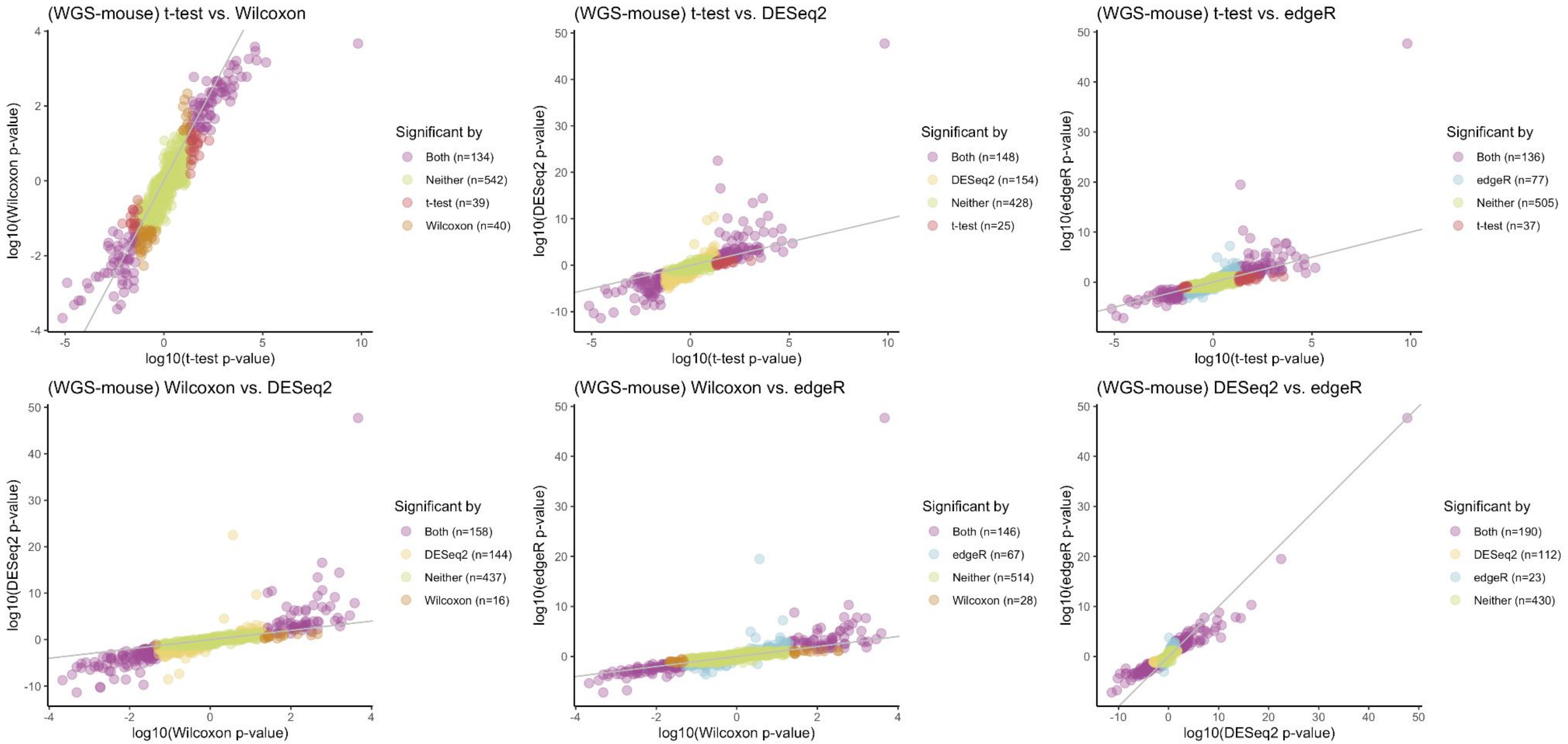
(WGS-mouse) t-test vs. Wilcoxon
log10(Wilcoxon p-value)
log10(t-test p-value)
Significant by
Both (n=134)
Neither (n=542)
t-test (n=39)
Wilcoxon (n=40)
(WGS-mouse) t-test vs. DESeq2
log10(DESeq2 p-value)
log10(t-test p-value)
Significant by
Both (n=148)
DESeq2 (n=154)
Neither (n=428)
t-test (n=25)
(WGS-mouse) t-test vs. edgeR
log10(edgeR p-value)
log10(t-test p-value)
Significant by
Both (n=136)
edgeR (n=77)
Neither (n=505)
t-test (n=37)
(WGS-mouse) Wilcoxon vs. DESeq2
log10(DESeq2 p-value)
log10(Wilcoxon p-value)
Significant by
Both (n=158)
DESeq2 (n=144)
Neither (n=437)
Wilcoxon (n=16)
(WGS-mouse) Wilcoxon vs. edgeR
log10(edgeR p-value)
log10(Wilcoxon p-value)
Significant by
Both (n=146)
edgeR (n=67)
Neither (n=514)
Wilcoxon (n=28)
(WGS-mouse) DESeq2 vs. edgeR
log10(edgeR p-value)
log10(DESeq2 p-value)
Significant by
Both (n=190)
DESeq2 (n=112)
edgeR (n=23)
Neither (n=430)

S7.

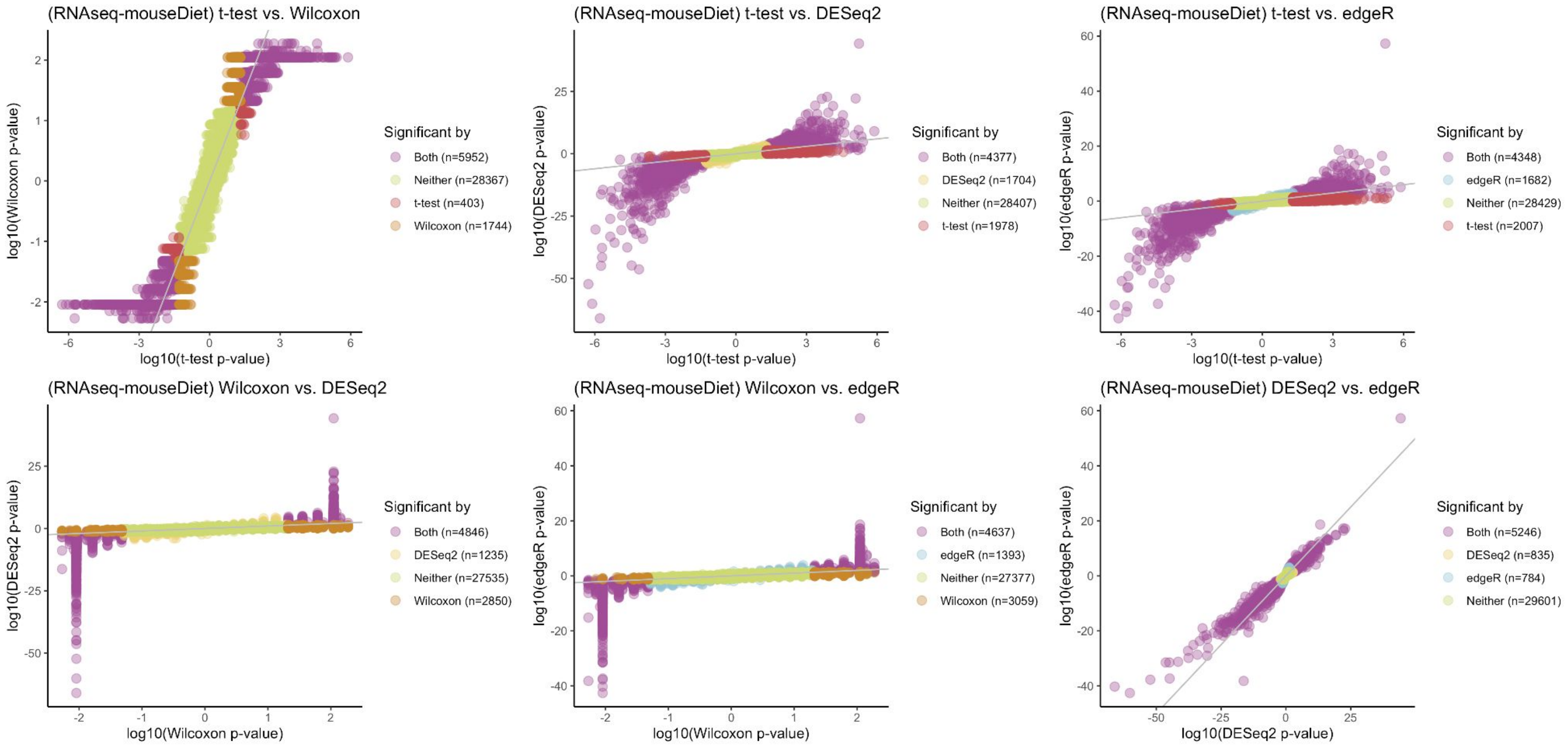
(RNAseq-mouseDiet) t-test vs. Wilcoxon
log10(Wilcoxon p-value)
log10(t-test p-value)
Significant by
Both (n=5952)
Neither (n=28367)
t-test (n=403)
Wilcoxon (n=1744)
(RNAseq-mouseDiet) t-test vs. DESeq2
log10(DESeq2 p-value)
log10(t-test p-value)
Significant by
Both (n=4377)
DESeq2 (n=1704)
Neither (n=28407)
t-test (n=1978)
(RNAseq-mouseDiet) t-test vs. edgeR
log10(edgeR p-value)
log10(t-test p-value)
Significant by
Both (n=4348)
edgeR (n=1682)
Neither (n=28429)
t-test (n=2007)
(RNAseq-mouseDiet) Wilcoxon vs. DESeq2
log10(DESeq2 p-value)
log10(Wilcoxon p-value)
Significant by
Both (n=4846)
DESeq2 (n=1235)
Neither (n=27535)
Wilcoxon (n=2850)
(RNAseq-mouseDiet) Wilcoxon vs. edgeR
log10(edgeR p-value)
log10(Wilcoxon p-value)
Significant by
Both (n=4637)
edgeR (n=1393)
Neither (n=27377)
Wilcoxon (n=3059)
(RNAseq-mouseDiet) DESeq2 vs. edgeR
log10(edgeR p-value)
log10(DESeq2 p-value)
Significant by
Both (n=5246)
DESeq2 (n=835)
edgeR (n=784)
Neither (n=29601)

S8.

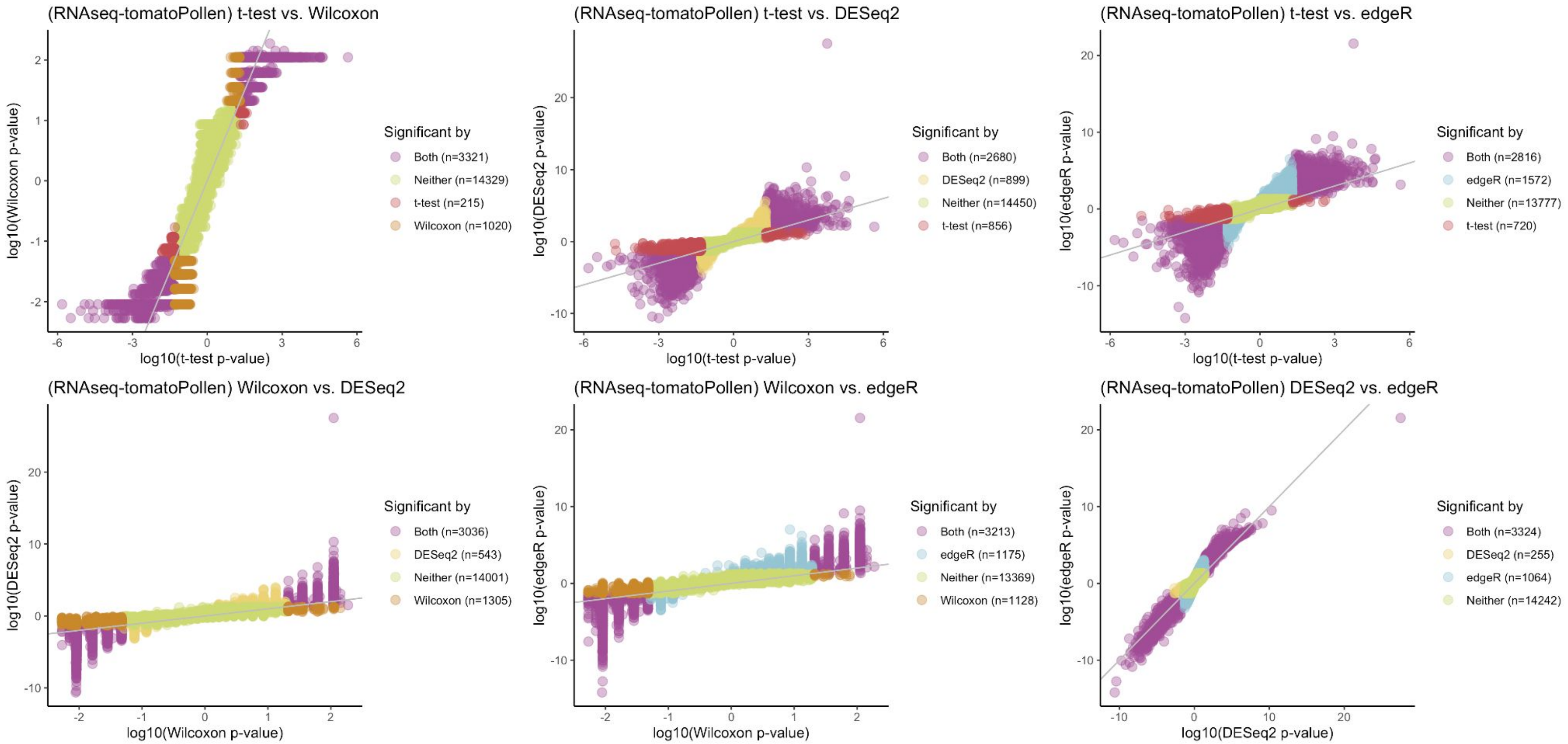
(RNAseq-tomatoPollen) t-test vs. Wilcoxon
log10(Wilcoxon p-value)
log10(t-test p-value)
Significant by
Both (n=3321)
Neither (n=14329)
t-test (n=215)
Wilcoxon (n=1020)
(RNAseq-tomatoPollen) t-test vs. DESeq2
log10(DESeq2 p-value)
log10(t-test p-value)
Significant by
Both (n=2680)
DESeq2 (n=899)
Neither (n=14450)
t-test (n=856)
(RNAseq-tomatoPollen) t-test vs. edgeR
log10(edgeR p-value)
log10(t-test p-value)
Significant by
Both (n=2816)
edgeR (n=1572)
Neither (n=13777)
t-test (n=720)
(RNAseq-tomatoPollen) Wilcoxon vs. DESeq2
log10(DESeq2 p-value)
log10(Wilcoxon p-value)
Significant by
Both (n=3036)
DESeq2 (n=543)
Neither (n=14001)
Wilcoxon (n=1305)
(RNAseq-tomatoPollen) Wilcoxon vs. edgeR
log10(edgeR p-value)
log10(Wilcoxon p-value)
Significant by
Both (n=3213)
edgeR (n=1175)
Neither (n=13369)
Wilcoxon (n=1128)
(RNAseq-tomatoPollen) DESeq2 vs. edgeR
log10(edgeR p-value)
log10(DESeq2 p-value)
Significant by
Both (n=3324)
DESeq2 (n=255)
edgeR (n=1064)
Neither (n=14242)

S9.

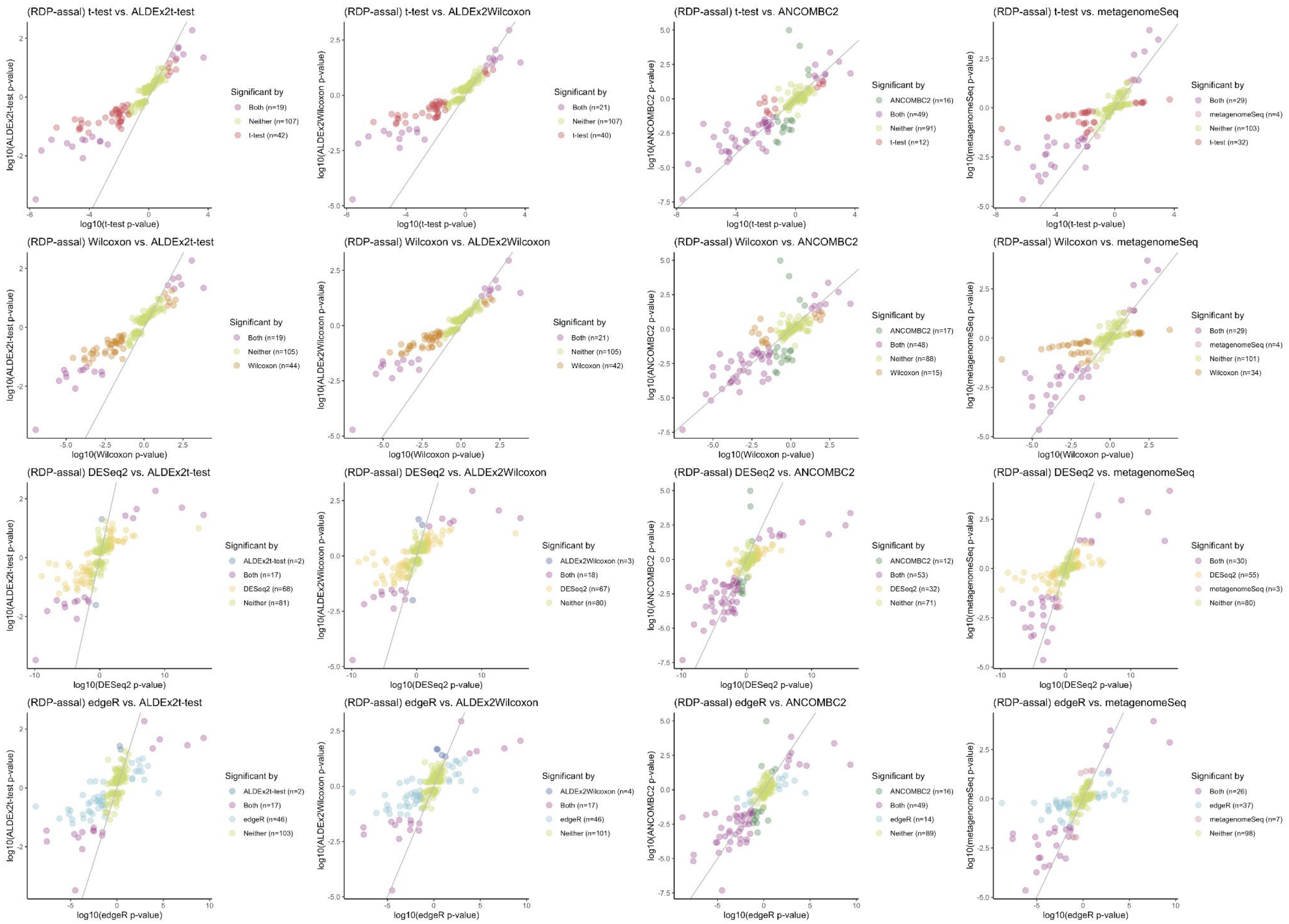
(RDP-assal) t-test vs. ALDEx2t-test
log10(ALDEx2t-test p-value)
log10(t-test p-value)
Significant by
Both (n=19)
Neither (n=107)
t-test (n=42)
(RDP-assal) t-test vs. ALDEx2Wilcoxon
log10(ALDEx2Wilcoxon p-value)
log10(t-test p-value)
Significant by
Both (n=21)
Neither (n=107)
t-test (n=40)
(RDP-assal) t-test vs. ANCOMBC2
log10(ANCOMBC2 p-value)
log10(t-test p-value)
Significant by
ANCOMBC2 (n=16)
Both (n=49)
Neither (n=91)
t-test (n=12)
(RDP-assal) t-test vs. metagenomeSeq
log10(metagenomeSeq p-value)
log10(t-test p-value)
Significant by
Both (n=29)
metagenomeSeq (n=4)
Neither (n=103)
t-test (n=32)
(RDP-assal) Wilcoxon vs. ALDEx2t-test
log10(ALDEx2t-test p-value)
log10(Wilcoxon p-value)
Significant by
Both (n=19)
Neither (n=105)
Wilcoxon (n=44)
(RDP-assal) Wilcoxon vs. ALDEx2Wilcoxon
log10(ALDEx2Wilcoxon p-value)
log10(Wilcoxon p-value)
Significant by
Both (n=21)
Neither (n=105)
Wilcoxon (n=42)
(RDP-assal) Wilcoxon vs. ANCOMBC2
log10(ANCOMBC2 p-value)
log10(Wilcoxon p-value)
Significant by
ANCOMBC2 (n=17)
Both (n=48)
Neither (n=88)
Wilcoxon (n=15)
(RDP-assal) Wilcoxon vs. metagenomeSeq
log10(metagenomeSeq p-value)
log10(Wilcoxon p-value)
Significant by
Both (n=29)
metagenomeSeq (n=4)
Neither (n=101)
Wilcoxon (n=34)
(RDP-assal) DESeq2 vs. ALDEx2t-test
log10(ALDEx2t-test p-value)
log10(DESeq2 p-value)
Significant by
ALDEx2t-test (n=2)
Both (n=17)
DESeq2 (n=68)
Neither (n=81)
(RDP-assal) DESeq2 vs. ALDEx2Wilcoxon
log10(ALDEx2Wilcoxon p-value)
log10(DESeq2 p-value)
Significant by
ALDEx2Wilcoxon (n=3)
Both (n=18)
DESeq2 (n=67)
Neither (n=80)
(RDP-assal) DESeq2 vs. ANCOMBC2
log10(ANCOMBC2 p-value)
log10(DESeq2 p-value)
Significant by
ANCOMBC2 (n=12)
Both (n=53)
DESeq2 (n=32)
Neither (n=71)
(RDP-assal) DESeq2 vs. metagenomeSeq
log10(metagenomeSeq p-value)
log10(DESeq2 p-value)
Significant by
Both (n=30)
DESeq2 (n=55)
metagenomeSeq (n=3)
Neither (n=80)
(RDP-assal) edgeR vs. ALDEx2t-test
log10(ALDEx2t-test p-value)
log10(edgeR p-value)
Significant by
ALDEx2t-test (n=2)
Both (n=17)
edgeR (n=46)
Neither (n=103)
(RDP-assal) edgeR vs. ALDEx2Wilcoxon
log10(ALDEx2Wilcoxon p-value)
log10(edgeR p-value)
Significant by
ALDEx2Wilcoxon (n=4)
Both (n=17)
edgeR (n=46)
Neither (n=101)
(RDP-assal) edgeR vs. ANCOMBC2
log10(ANCOMBC2 p-value)
log10(edgeR p-value)
Significant by
ANCOMBC2 (n=16)
Both (n=49)
edgeR (n=14)
Neither (n=89)
(RDP-assal) edgeR vs. metagenomeSeq
log10(metagenomeSeq p-value)
log10(edgeR p-value)
Significant by
Both (n=26)
edgeR (n=37)
metagenomeSeq (n=7)
Neither (n=98)

S10.

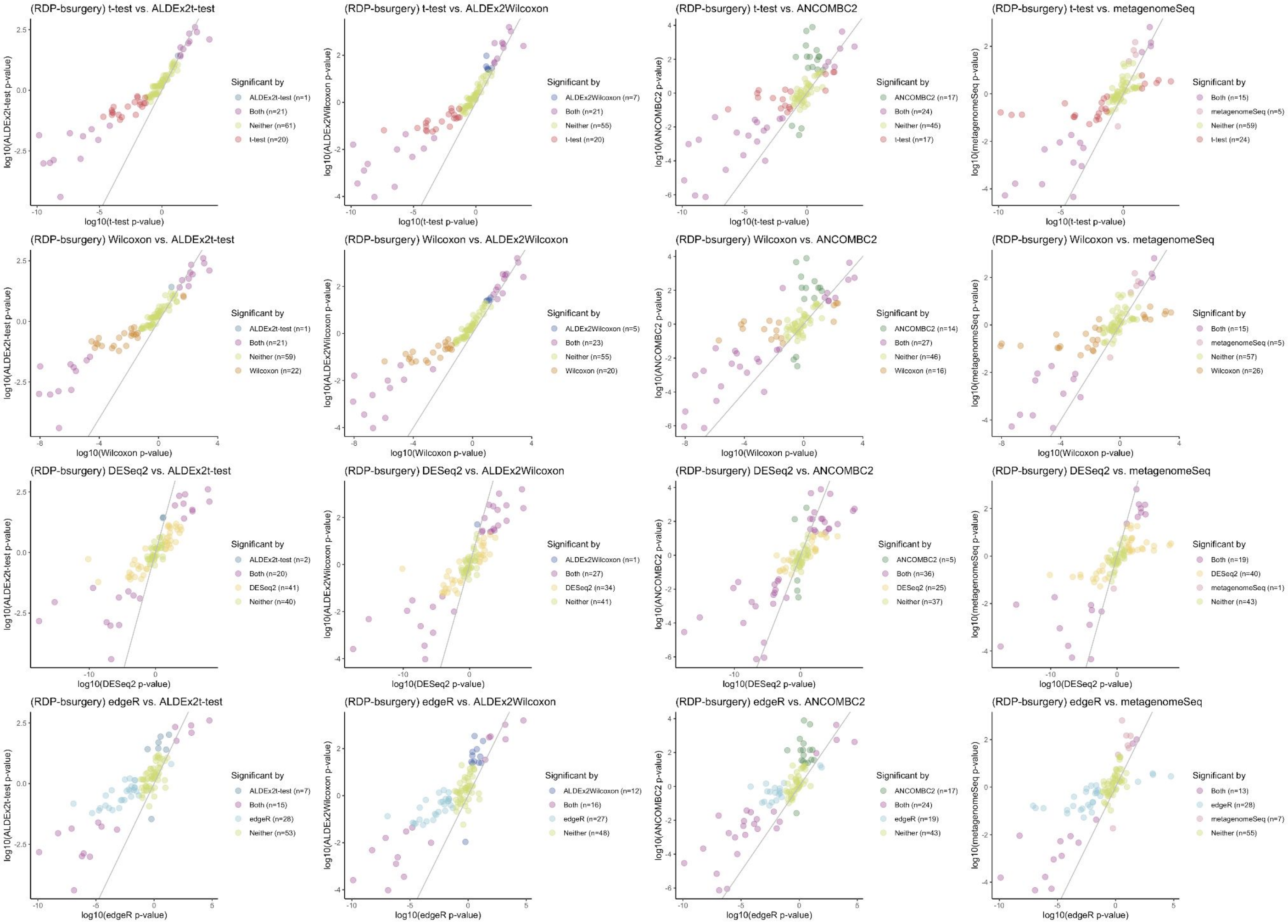
(RDP-bsurgery) t-test vs. ALDEx2t-test
log10(ALDEx2t-test p-value)
log10(t-test p-value)
Significant by
ALDEx2t-test (n=1)
Both (n=21)
Neither (n=61)
t-test (n=20)
(RDP-bsurgery) t-test vs. ALDEx2Wilcoxon
log10(ALDEx2Wilcoxon p-value)
log10(t-test p-value)
Significant by
ALDEx2Wilcoxon (n=7)
Both (n=21)
Neither (n=55)
t-test (n=20)
(RDP-bsurgery) t-test vs. ANCOMBC2
log10(ANCOMBC2 p-value)
log10(t-test p-value)
Significant by
ANCOMBC2 (n=17)
Both (n=24)
Neither (n=45)
t-test (n=17)
(RDP-bsurgery) t-test vs. metagenomeSeq
log10(metagenomeSeq p-value)
log10(t-test p-value)
Significant by
Both (n=15)
metagenomeSeq (n=5)
Neither (n=59)
t-test (n=24)
(RDP-bsurgery) Wilcoxon vs. ALDEx2t-test
log10(ALDEx2t-test p-value)
log10(Wilcoxon p-value)
Significant by
ALDEx2t-test (n=1)
Both (n=21)
Neither (n=59)
Wilcoxon (n=22)
(RDP-bsurgery) Wilcoxon vs. ALDEx2Wilcoxon
log10(ALDEx2Wilcoxon p-value)
log10(Wilcoxon p-value)
Significant by
ALDEx2Wilcoxon (n=5)
Both (n=23)
Neither (n=55)
Wilcoxon (n=20)
(RDP-bsurgery) Wilcoxon vs. ANCOMBC2
log10(ANCOMBC2 p-value)
log10(Wilcoxon p-value)
Significant by
ANCOMBC2 (n=14)
Both (n=27)
Neither (n=46)
Wilcoxon (n=16)
(RDP-bsurgery) Wilcoxon vs. metagenomeSeq
log10(metagenomeSeq p-value)
log10(Wilcoxon p-value)
Significant by
Both (n=15)
metagenomeSeq (n=5)
Neither (n=57)
Wilcoxon (n=26)
(RDP-bsurgery) DESeq2 vs. ALDEx2t-test
log10(ALDEx2t-test p-value)
log10(DESeq2 p-value)
Significant by
ALDEx2t-test (n=2)
Both (n=20)
DESeq2 (n=41)
Neither (n=40)
(RDP-bsurgery) DESeq2 vs. ALDEx2Wilcoxon
log10(ALDEx2Wilcoxon p-value)
log10(DESeq2 p-value)
Significant by
ALDEx2Wilcoxon (n=1)
Both (n=27)
DESeq2 (n=34)
Neither (n=41)
(RDP-bsurgery) DESeq2 vs. ANCOMBC2
log10(ANCOMBC2 p-value)
log10(DESeq2 p-value)
Significant by
ANCOMBC2 (n=5)
Both (n=36)
DESeq2 (n=25)
Neither (n=37)
(RDP-bsurgery) DESeq2 vs. metagenomeSeq
log10(metagenomeSeq p-value)
log10(DESeq2 p-value)
Significant by
Both (n=19)
DESeq2 (n=40)
metagenomeSeq (n=1)
Neither (n=43)
(RDP-bsurgery) edgeR vs. ALDEx2t-test
log10(ALDEx2t-test p-value)
log10(edgeR p-value)
Significant by
ALDEx2t-test (n=7)
Both (n=15)
edgeR (n=28)
Neither (n=53)
(RDP-bsurgery) edgeR vs. ALDEx2Wilcoxon
log10(ALDEx2Wilcoxon p-value)
log10(edgeR p-value)
Significant by
ALDEx2Wilcoxon (n=12)
Both (n=16)
edgeR (n=27)
Neither (n=48)
(RDP-bsurgery) edgeR vs. ANCOMBC2
log10(ANCOMBC2 p-value)
log10(edgeR p-value)
Significant by
ANCOMBC2 (n=17)
Both (n=24)
edgeR (n=19)
Neither (n=43)
(RDP-bsurgery) edgeR vs. metagenomeSeq
log10(metagenomeSeq p-value)
log10(edgeR p-value)
Significant by
Both (n=13)
edgeR (n=28)
metagenomeSeq (n=7)
Neither (n=55)

S11.

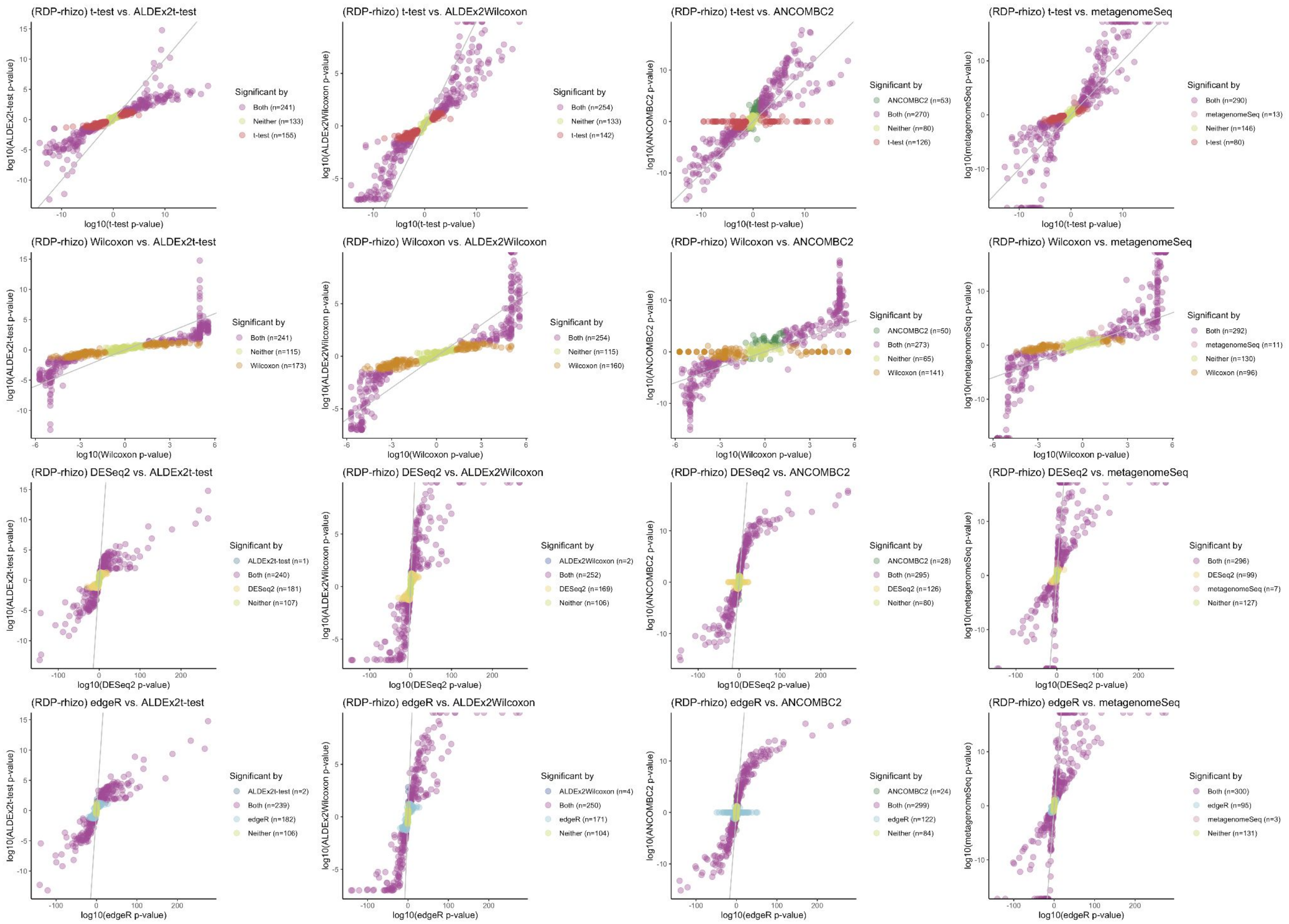
(RDP-rhizo) t-test vs. ALDEx2t-test
log10(ALDEx2t-test p-value)
log10(t-test p-value)
Significant by
Both (n=241)
Neither (n=133)
t-test (n=155)
(RDP-rhizo) t-test vs. ALDEx2Wilcoxon
log10(ALDEx2Wilcoxon p-value)
log10(t-test p-value)
Significant by
Both (n=254)
Neither (n=133)
t-test (n=142)
(RDP-rhizo) t-test vs. ANCOMBC2
log10(ANCOMBC2 p-value)
log10(t-test p-value)
Significant by
ANCOMBC2 (n=53)
Both (n=270)
Neither (n=80)
t-test (n=126)
(RDP-rhizo) t-test vs. metagenomeSeq
log10(metagenomeSeq p-value)
log10(t-test p-value)
Significant by
Both (n=290)
metagenomeSeq (n=13)
Neither (n=146)
t-test (n=80)
(RDP-rhizo) Wilcoxon vs. ALDEx2t-test
log10(ALDEx2t-test p-value)
log10(Wilcoxon p-value)
Significant by
Both (n=241)
Neither (n=115)
Wilcoxon (n=173)
(RDP-rhizo) Wilcoxon vs. ALDEx2Wilcoxon
log10(ALDEx2Wilcoxon p-value)
log10(Wilcoxon p-value)
Significant by
Both (n=254)
Neither (n=115)
Wilcoxon (n=160)
(RDP-rhizo) Wilcoxon vs. ANCOMBC2
log10(ANCOMBC2 p-value)
log10(Wilcoxon p-value)
Significant by
ANCOMBC2 (n=50)
Both (n=273)
Neither (n=65)
Wilcoxon (n=141)
(RDP-rhizo) Wilcoxon vs. metagenomeSeq
log10(metagenomeSeq p-value)
log10(Wilcoxon p-value)
Significant by
Both (n=292)
metagenomeSeq (n=11)
Neither (n=130)
Wilcoxon (n=96)
(RDP-rhizo) DESeq2 vs. ALDEx2t-test
log10(ALDEx2t-test p-value)
log10(DESeq2 p-value)
Significant by
ALDEx2t-test (n=1)
Both (n=240)
DESeq2 (n=181)
Neither (n=107)
(RDP-rhizo) DESeq2 vs. ALDEx2Wilcoxon
log10(ALDEx2Wilcoxon p-value)
log10(DESeq2 p-value)
Significant by
ALDEx2Wilcoxon (n=2)
Both (n=252)
DESeq2 (n=169)
Neither (n=106)
(RDP-rhizo) DESeq2 vs. ANCOMBC2
log10(ANCOMBC2 p-value)
log10(DESeq2 p-value)
Significant by
ANCOMBC2 (n=28)
Both (n=295)
DESeq2 (n=126)
Neither (n=80)
(RDP-rhizo) DESeq2 vs. metagenomeSeq
log10(metagenomeSeq p-value)
log10(DESeq2 p-value)
Significant by
Both (n=296)
DESeq2 (n=99)
metagenomeSeq (n=7)
Neither (n=127)
(RDP-rhizo) edgeR vs. ALDEx2t-test
log10(ALDEx2t-test p-value)
log10(edgeR p-value)
Significant by
ALDEx2t-test (n=2)
Both (n=239)
edgeR (n=182)
Neither (n=106)
(RDP-rhizo) edgeR vs. ALDEx2Wilcoxon
log10(ALDEx2Wilcoxon p-value)
log10(edgeR p-value)
Significant by
ALDEx2Wilcoxon (n=4)
Both (n=250)
edgeR (n=171)
Neither (n=104)
(RDP-rhizo) edgeR vs. ANCOMBC2
log10(ANCOMBC2 p-value)
log10(edgeR p-value)
Significant by
ANCOMBC2 (n=24)
Both (n=299)
edgeR (n=122)
Neither (n=84)
(RDP-rhizo) edgeR vs. metagenomeSeq
log10(metagenomeSeq p-value)
log10(edgeR p-value)
Significant by
Both (n=300)
edgeR (n=95)
metagenomeSeq (n=3)
Neither (n=131)

S12.

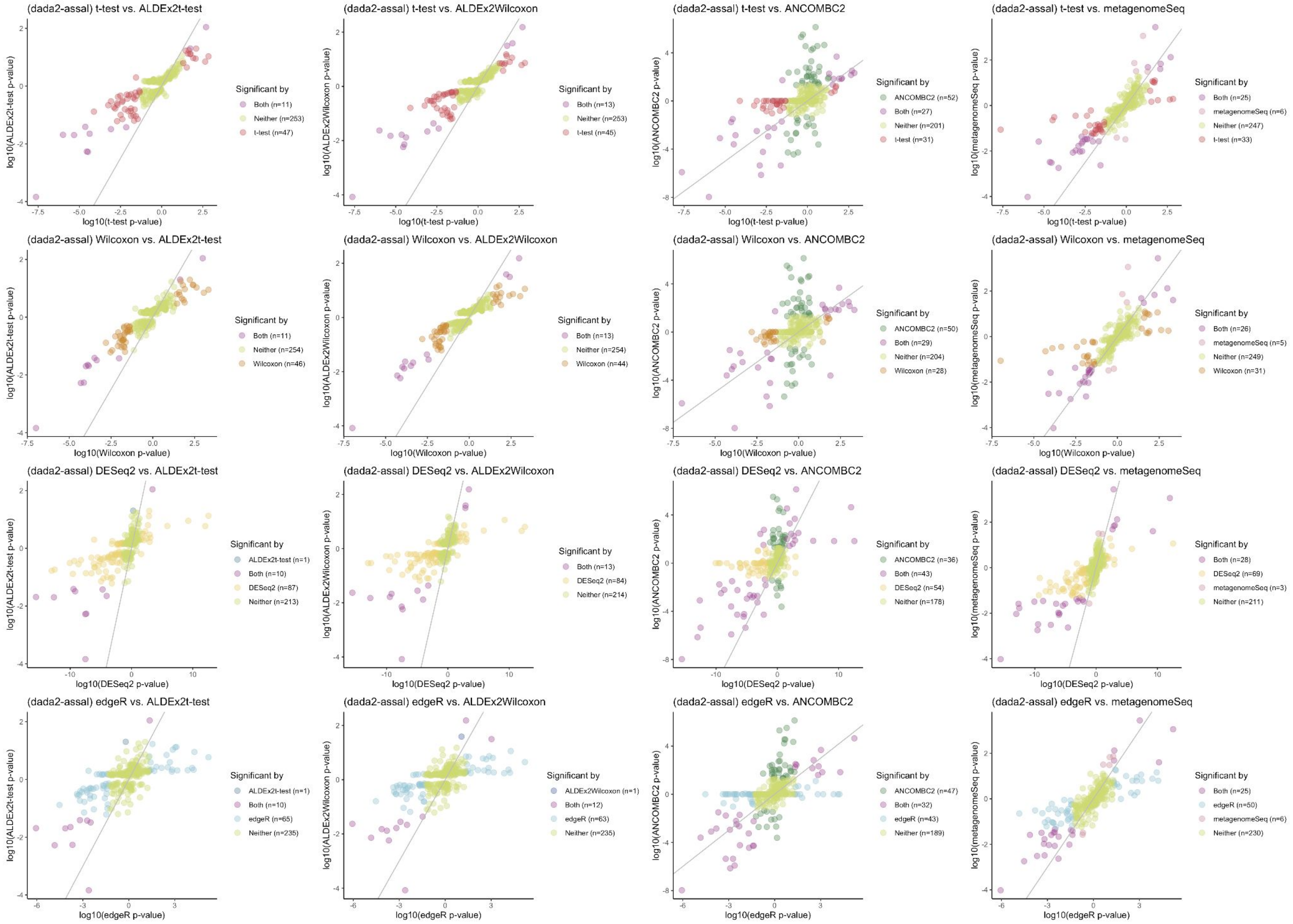
(dada2-assal) t-test vs. ALDEx2t-test
log10(ALDEx2t-test p-value)
log10(t-test p-value)
Significant by
Both (n=11)
Neither (n=253)
t-test (n=47)
(dada2-assal) t-test vs. ALDEx2Wilcoxon
log10(ALDEx2Wilcoxon p-value)
log10(t-test p-value)
Significant by
Both (n=13)
Neither (n=253)
t-test (n=45)
(dada2-assal) t-test vs. ANCOMBC2
log10(ANCOMBC2 p-value)
log10(t-test p-value)
Significant by
ANCOMBC2 (n=52)
Both (n=27)
Neither (n=201)
t-test (n=31)
(dada2-assal) t-test vs. metagenomeSeq
log10(metagenomeSeq p-value)
log10(t-test p-value)
Significant by
Both (n=25)
metagenomeSeq (n=6)
Neither (n=247)
t-test (n=33)
(dada2-assal) Wilcoxon vs. ALDEx2t-test
log10(ALDEx2t-test p-value)
log10(Wilcoxon p-value)
Significant by
Both (n=11)
Neither (n=254)
Wilcoxon (n=46)
(dada2-assal) Wilcoxon vs. ALDEx2Wilcoxon
log10(ALDEx2Wilcoxon p-value)
log10(Wilcoxon p-value)
Significant by
Both (n=13)
Neither (n=254)
Wilcoxon (n=44)
(dada2-assal) Wilcoxon vs. ANCOMBC2
log10(ANCOMBC2 p-value)
log10(Wilcoxon p-value)
Significant by
ANCOMBC2 (n=50)
Both (n=29)
Neither (n=204)
Wilcoxon (n=28)
(dada2-assal) Wilcoxon vs. metagenomeSeq
log10(metagenomeSeq p-value)
log10(Wilcoxon p-value)
Significant by
Both (n=26)
metagenomeSeq (n=5)
Neither (n=249)
Wilcoxon (n=31)
(dada2-assal) DESeq2 vs. ALDEx2t-test
log10(ALDEx2t-test p-value)
log10(DESeq2 p-value)
Significant by
ALDEx2t-test (n=1)
Both (n=10)
DESeq2 (n=87)
Neither (n=213)
(dada2-assal) DESeq2 vs. ALDEx2Wilcoxon
log10(ALDEx2Wilcoxon p-value)
log10(DESeq2 p-value)
Significant by
Both (n=13)
DESeq2 (n=84)
Neither (n=214)
(dada2-assal) DESeq2 vs. ANCOMBC2
log10(ANCOMBC2 p-value)
log10(DESeq2 p-value)
Significant by
ANCOMBC2 (n=36)
Both (n=43)
DESeq2 (n=54)
Neither (n=178)
(dada2-assal) DESeq2 vs. metagenomeSeq
log10(metagenomeSeq p-value)
log10(DESeq2 p-value)
Significant by
Both (n=28)
DESeq2 (n=69)
metagenomeSeq (n=3)
Neither (n=211)
(dada2-assal) edgeR vs. ALDEx2t-test
log10(ALDEx2t-test p-value)
log10(edgeR p-value)
Significant by
ALDEx2t-test (n=1)
Both (n=10)
edgeR (n=65)
Neither (n=235)
(dada2-assal) edgeR vs. ALDEx2Wilcoxon
log10(ALDEx2Wilcoxon p-value)
log10(edgeR p-value)
Significant by
ALDEx2Wilcoxon (n=1)
Both (n=12)
edgeR (n=63)
Neither (n=235)
(dada2-assal) edgeR vs. ANCOMBC2
log10(ANCOMBC2 p-value)
log10(edgeR p-value)
Significant by
ANCOMBC2 (n=47)
Both (n=32)
edgeR (n=43)
Neither (n=189)
(dada2-assal) edgeR vs. metagenomeSeq
log10(metagenomeSeq p-value)
log10(edgeR p-value)
Significant by
Both (n=25)
edgeR (n=50)
metagenomeSeq (n=6)
Neither (n=230)

S13.

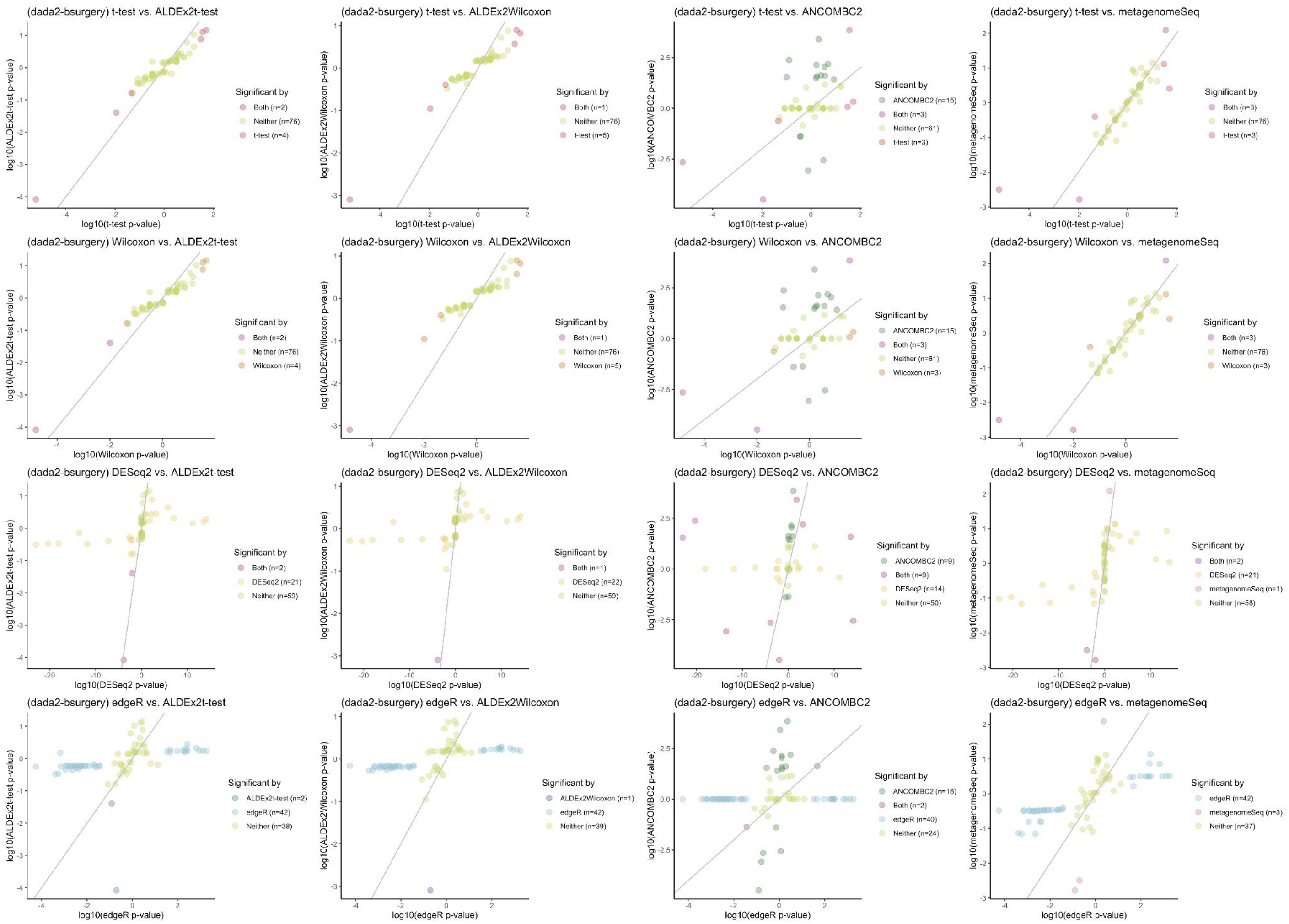
(dada2-bsurgery) t-test vs. ALDEx2t-test
log10(t-test p-value)
log10(ALDEx2t-test p-value)
Significant by
Both (n=2)
Neither (n=76)
t-test (n=4)
(dada2-bsurgery) t-test vs. ALDEx2Wilcoxon
log10(t-test p-value)
log10(ALDEx2Wilcoxon p-value)
Significant by
Both (n=1)
Neither (n=76)
t-test (n=5)
(dada2-bsurgery) t-test vs. ANCOMBC2
log10(t-test p-value)
log10(ANCOMBC2 p-value)
Significant by
ANCOMBC2 (n=15)
Both (n=3)
Neither (n=61)
t-test (n=3)
(dada2-bsurgery) t-test vs. metagenomeSeq
log10(t-test p-value)
log10(metagenomeSeq p-value)
Significant by
Both (n=3)
Neither (n=76)
t-test (n=3)
(dada2-bsurgery) Wilcoxon vs. ALDEx2t-test
log10(Wilcoxon p-value)
log10(ALDEx2t-test p-value)
Significant by
Both (n=2)
Neither (n=76)
Wilcoxon (n=4)
(dada2-bsurgery) Wilcoxon vs. ALDEx2Wilcoxon
log10(Wilcoxon p-value)
log10(ALDEx2Wilcoxon p-value)
Significant by
Both (n=1)
Neither (n=76)
Wilcoxon (n=5)
(dada2-bsurgery) Wilcoxon vs. ANCOMBC2
log10(Wilcoxon p-value)
log10(ANCOMBC2 p-value)
Significant by
ANCOMBC2 (n=15)
Both (n=3)
Neither (n=61)
Wilcoxon (n=3)
(dada2-bsurgery) Wilcoxon vs. metagenomeSeq
log10(Wilcoxon p-value)
log10(metagenomeSeq p-value)
Significant by
Both (n=3)
Neither (n=76)
Wilcoxon (n=3)
(dada2-bsurgery) DESeq2 vs. ALDEx2t-test
log10(DESeq2 p-value)
log10(ALDEx2t-test p-value)
Significant by
Both (n=2)
DESeq2 (n=21)
Neither (n=59)
(dada2-bsurgery) DESeq2 vs. ALDEx2Wilcoxon
log10(DESeq2 p-value)
log10(ALDEx2Wilcoxon p-value)
Significant by
Both (n=1)
DESeq2 (n=22)
Neither (n=59)
(dada2-bsurgery) DESeq2 vs. ANCOMBC2
log10(DESeq2 p-value)
log10(ANCOMBC2 p-value)
Significant by
ANCOMBC2 (n=9)
Both (n=9)
DESeq2 (n=14)
Neither (n=50)
(dada2-bsurgery) DESeq2 vs. metagenomeSeq
log10(DESeq2 p-value)
log10(metagenomeSeq p-value)
Significant by
Both (n=2)
DESeq2 (n=21)
metagenomeSeq (n=1)
Neither (n=58)
(dada2-bsurgery) edgeR vs. ALDEx2t-test
log10(edgeR p-value)
log10(ALDEx2t-test p-value)
Significant by
ALDEx2t-test (n=2)
edgeR (n=42)
Neither (n=38)
(dada2-bsurgery) edgeR vs. ALDEx2Wilcoxon
log10(edgeR p-value)
log10(ALDEx2Wilcoxon p-value)
Significant by
ALDEx2Wilcoxon (n=1)
edgeR (n=42)
Neither (n=39)
(dada2-bsurgery) edgeR vs. ANCOMBC2
log10(edgeR p-value)
log10(ANCOMBC2 p-value)
Significant by
ANCOMBC2 (n=16)
Both (n=2)
edgeR (n=40)
Neither (n=24)
(dada2-bsurgery) edgeR vs. metagenomeSeq
log10(edgeR p-value)
log10(metagenomeSeq p-value)
Significant by
edgeR (n=42)
metagenomeSeq (n=3)
Neither (n=37)

S14.

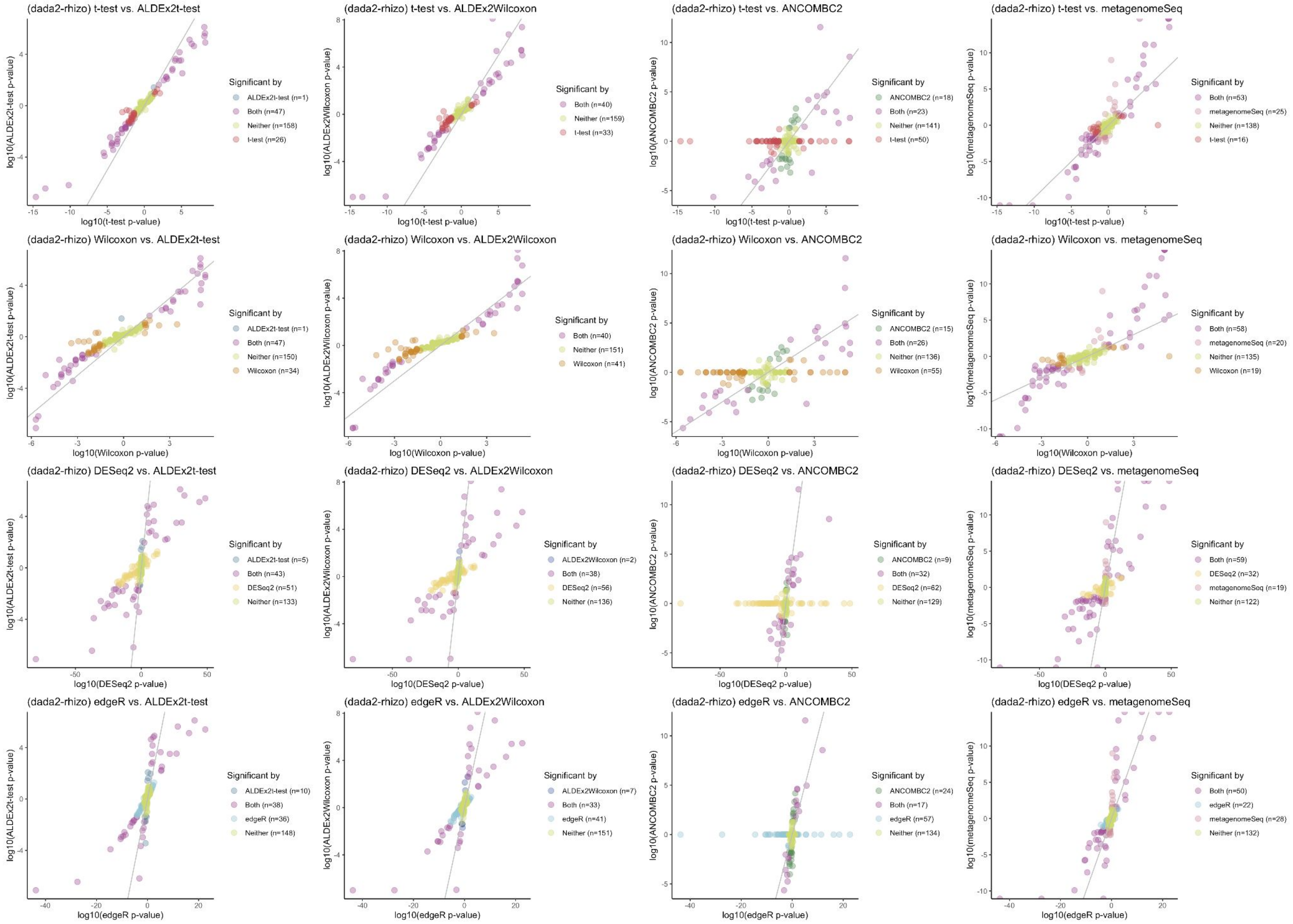
(dada2-rhizo) t-test vs. ALDEx2t-test
log10(ALDEx2t-test p-value)
log10(t-test p-value)
Significant by
ALDEx2t-test (n=1)
Both (n=47)
Neither (n=158)
t-test (n=26)
(dada2-rhizo) t-test vs. ALDEx2Wilcoxon
log10(ALDEx2Wilcoxon p-value)
log10(t-test p-value)
Significant by
Both (n=40)
Neither (n=159)
t-test (n=33)
(dada2-rhizo) t-test vs. ANCOMBC2
log10(ANCOMBC2 p-value)
log10(t-test p-value)
Significant by
ANCOMBC2 (n=18)
Both (n=23)
Neither (n=141)
t-test (n=50)
(dada2-rhizo) t-test vs. metagenomeSeq
log10(metagenomeSeq p-value)
log10(t-test p-value)
Significant by
Both (n=53)
metagenomeSeq (n=25)
Neither (n=138)
t-test (n=16)
(dada2-rhizo) Wilcoxon vs. ALDEx2t-test
log10(ALDEx2t-test p-value)
log10(Wilcoxon p-value)
Significant by
ALDEx2t-test (n=1)
Both (n=47)
Neither (n=150)
Wilcoxon (n=34)
(dada2-rhizo) Wilcoxon vs. ALDEx2Wilcoxon
log10(ALDEx2Wilcoxon p-value)
log10(Wilcoxon p-value)
Significant by
Both (n=40)
Neither (n=151)
Wilcoxon (n=41)
(dada2-rhizo) Wilcoxon vs. ANCOMBC2
log10(ANCOMBC2 p-value)
log10(Wilcoxon p-value)
Significant by
ANCOMBC2 (n=15)
Both (n=26)
Neither (n=136)
Wilcoxon (n=55)
(dada2-rhizo) Wilcoxon vs. metagenomeSeq
log10(metagenomeSeq p-value)
log10(Wilcoxon p-value)
Significant by
Both (n=58)
metagenomeSeq (n=20)
Neither (n=135)
Wilcoxon (n=19)
(dada2-rhizo) DESeq2 vs. ALDEx2t-test
log10(ALDEx2t-test p-value)
log10(DESeq2 p-value)
Significant by
ALDEx2t-test (n=5)
Both (n=43)
DESeq2 (n=51)
Neither (n=133)
(dada2-rhizo) DESeq2 vs. ALDEx2Wilcoxon
log10(ALDEx2Wilcoxon p-value)
log10(DESeq2 p-value)
Significant by
ALDEx2Wilcoxon (n=2)
Both (n=38)
DESeq2 (n=56)
Neither (n=136)
(dada2-rhizo) DESeq2 vs. ANCOMBC2
log10(ANCOMBC2 p-value)
log10(DESeq2 p-value)
Significant by
ANCOMBC2 (n=9)
Both (n=32)
DESeq2 (n=62)
Neither (n=129)
(dada2-rhizo) DESeq2 vs. metagenomeSeq
log10(metagenomeSeq p-value)
log10(DESeq2 p-value)
Significant by
Both (n=59)
DESeq2 (n=32)
metagenomeSeq (n=19)
Neither (n=122)
(dada2-rhizo) edgeR vs. ALDEx2t-test
log10(ALDEx2t-test p-value)
log10(edgeR p-value)
Significant by
ALDEx2t-test (n=10)
Both (n=38)
edgeR (n=36)
Neither (n=148)
(dada2-rhizo) edgeR vs. ALDEx2Wilcoxon
log10(ALDEx2Wilcoxon p-value)
log10(edgeR p-value)
Significant by
ALDEx2Wilcoxon (n=7)
Both (n=33)
edgeR (n=41)
Neither (n=151)
(dada2-rhizo) edgeR vs. ANCOMBC2
log10(ANCOMBC2 p-value)
log10(edgeR p-value)
Significant by
ANCOMBC2 (n=24)
Both (n=17)
edgeR (n=57)
Neither (n=134)
(dada2-rhizo) edgeR vs. metagenomeSeq
log10(metagenomeSeq p-value)
log10(edgeR p-value)
Significant by
Both (n=50)
edgeR (n=22)
metagenomeSeq (n=28)
Neither (n=132)

S15.

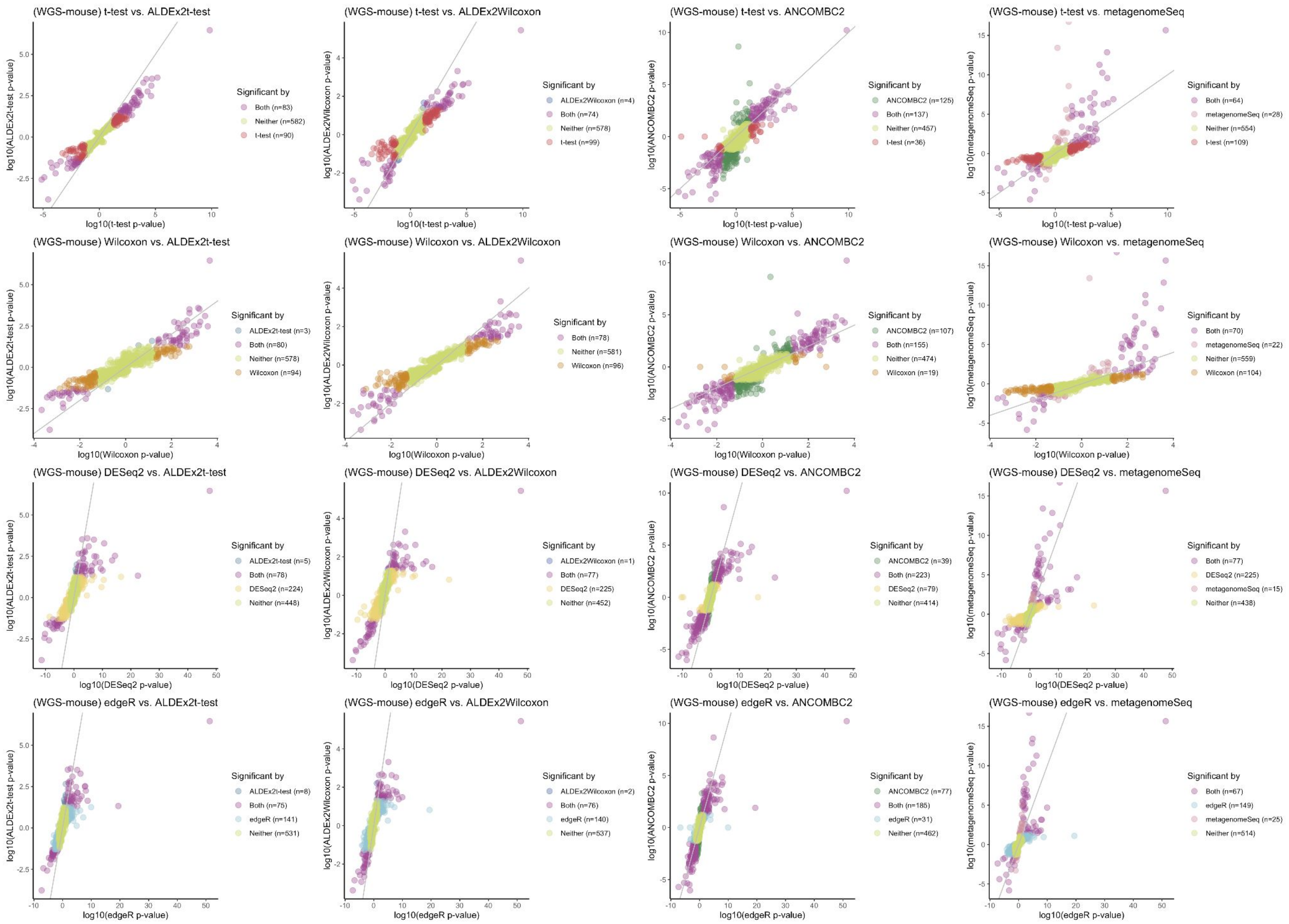

(WGS-mouse) t-test vs. ALDEx2t-test
log10(ALDEx2t-test p-value)
log10(t-test p-value)
Significant by
Both (n=83)
Neither (n=582)
t-test (n=90)
(WGS-mouse) t-test vs. ALDEx2Wilcoxon
log10(ALDEx2Wilcoxon p-value)
log10(t-test p-value)
Significant by
ALDEx2Wilcoxon (n=4)
Both (n=74)
Neither (n=578)
t-test (n=99)
(WGS-mouse) t-test vs. ANCOMBC2
log10(ANCOMBC2 p-value)
log10(t-test p-value)
Significant by
ANCOMBC2 (n=125)
Both (n=137)
Neither (n=457)
t-test (n=36)
(WGS-mouse) t-test vs. metagenomeSeq
log10(metagenomeSeq p-value)
log10(t-test p-value)
Significant by
Both (n=64)
metagenomeSeq (n=28)
Neither (n=554)
t-test (n=109)
(WGS-mouse) Wilcoxon vs. ALDEx2t-test
log10(ALDEx2t-test p-value)
log10(Wilcoxon p-value)
Significant by
ALDEx2t-test (n=3)
Both (n=80)
Neither (n=578)
Wilcoxon (n=94)
(WGS-mouse) Wilcoxon vs. ALDEx2Wilcoxon
log10(ALDEx2Wilcoxon p-value)
log10(Wilcoxon p-value)
Significant by
Both (n=78)
Neither (n=581)
Wilcoxon (n=96)
(WGS-mouse) Wilcoxon vs. ANCOMBC2
log10(ANCOMBC2 p-value)
log10(Wilcoxon p-value)
Significant by
ANCOMBC2 (n=107)
Both (n=155)
Neither (n=474)
Wilcoxon (n=19)
(WGS-mouse) Wilcoxon vs. metagenomeSeq
log10(metagenomeSeq p-value)
log10(Wilcoxon p-value)
Significant by
Both (n=70)
metagenomeSeq (n=22)
Neither (n=559)
Wilcoxon (n=104)
(WGS-mouse) DESeq2 vs. ALDEx2t-test
log10(ALDEx2t-test p-value)
log10(DESeq2 p-value)
Significant by
ALDEx2t-test (n=5)
Both (n=78)
DESeq2 (n=224)
Neither (n=448)
(WGS-mouse) DESeq2 vs. ALDEx2Wilcoxon
log10(ALDEx2Wilcoxon p-value)
log10(DESeq2 p-value)
Significant by
ALDEx2Wilcoxon (n=1)
Both (n=77)
DESeq2 (n=225)
Neither (n=452)
(WGS-mouse) DESeq2 vs. ANCOMBC2
log10(ANCOMBC2 p-value)
log10(DESeq2 p-value)
Significant by
ANCOMBC2 (n=39)
Both (n=223)
DESeq2 (n=79)
Neither (n=414)
(WGS-mouse) DESeq2 vs. metagenomeSeq
log10(metagenomeSeq p-value)
log10(DESeq2 p-value)
Significant by
Both (n=77)
DESeq2 (n=225)
metagenomeSeq (n=15)
Neither (n=438)
(WGS-mouse) edgeR vs. ALDEx2t-test
log10(ALDEx2t-test p-value)
log10(edgeR p-value)
Significant by
ALDEx2t-test (n=8)
Both (n=75)
edgeR (n=141)
Neither (n=531)
(WGS-mouse) edgeR vs. ALDEx2Wilcoxon
log10(ALDEx2Wilcoxon p-value)
log10(edgeR p-value)
Significant by
ALDEx2Wilcoxon (n=2)
Both (n=76)
edgeR (n=140)
Neither (n=537)
(WGS-mouse) edgeR vs. ANCOMBC2
log10(ANCOMBC2 p-value)
log10(edgeR p-value)
Significant by
ANCOMBC2 (n=77)
Both (n=185)
edgeR (n=31)
Neither (n=462)
(WGS-mouse) edgeR vs. metagenomeSeq
log10(metagenomeSeq p-value)
log10(edgeR p-value)
Significant by
Both (n=67)
edgeR (n=149)
metagenomeSeq (n=25)
Neither (n=514)

S16.

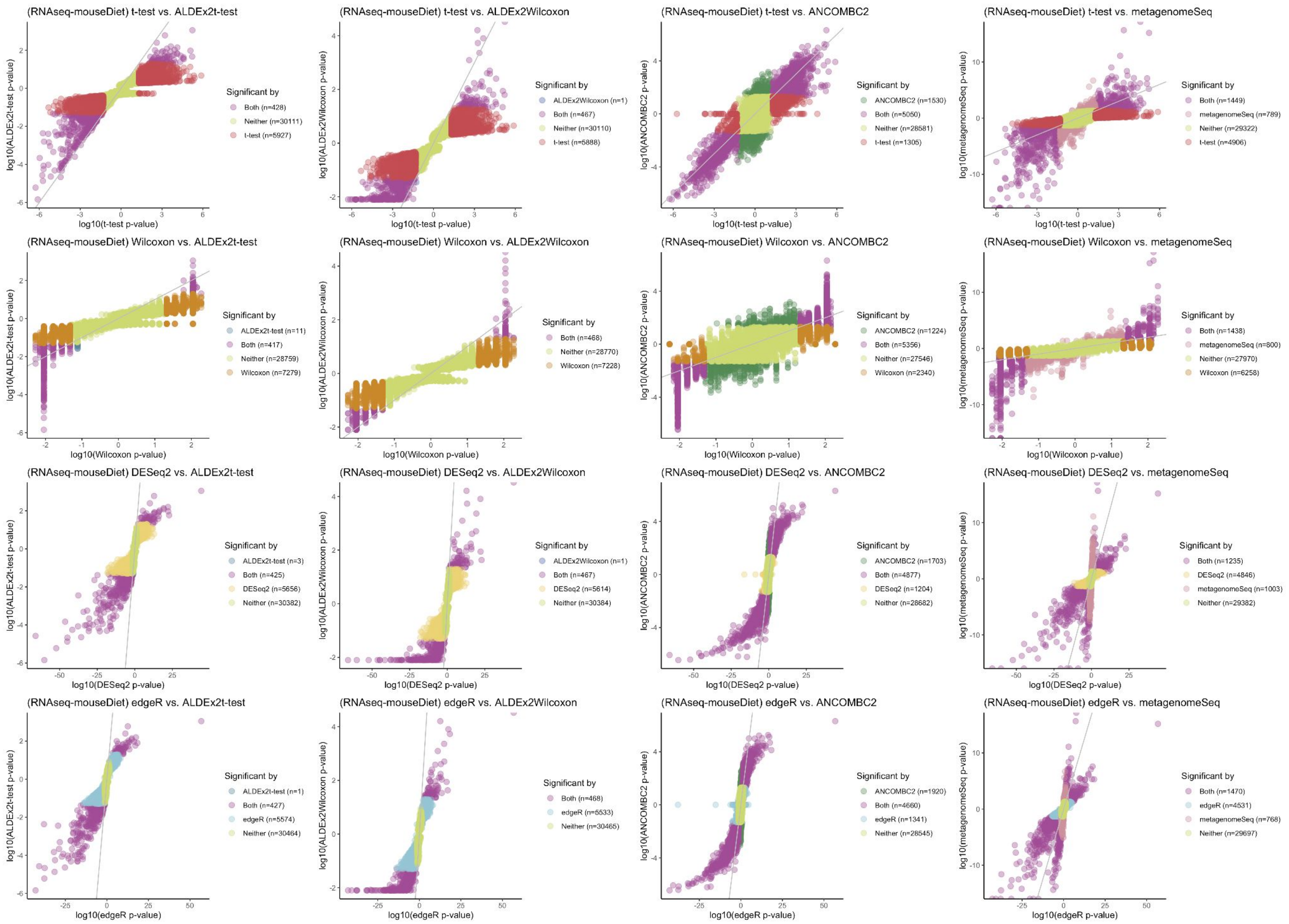
(RNAseq-mouseDiet) t-test vs. ALDEx2t-test
log10(ALDEx2t-test p-value)
log10(t-test p-value)
Significant by
Both (n=428)
Neither (n=30111)
t-test (n=5927)
(RNAseq-mouseDiet) t-test vs. ALDEx2Wilcoxon
log10(ALDEx2Wilcoxon p-value)
log10(t-test p-value)
Significant by
ALDEx2Wilcoxon (n=1)
Both (n=467)
Neither (n=30110)
t-test (n=5888)
(RNAseq-mouseDiet) t-test vs. ANCOMBC2
log10(ANCOMBC2 p-value)
log10(t-test p-value)
Significant by
ANCOMBC2 (n=1530)
Both (n=5050)
Neither (n=28581)
t-test (n=1305)
(RNAseq-mouseDiet) t-test vs. metagenomeSeq
log10(metagenomeSeq p-value)
log10(t-test p-value)
Significant by
Both (n=1449)
metagenomeSeq (n=789)
Neither (n=29322)
t-test (n=4906)
(RNAseq-mouseDiet) Wilcoxon vs. ALDEx2t-test
log10(ALDEx2t-test p-value)
log10(Wilcoxon p-value)
Significant by
ALDEx2t-test (n=11)
Both (n=417)
Neither (n=28759)
Wilcoxon (n=7279)
(RNAseq-mouseDiet) Wilcoxon vs. ALDEx2Wilcoxon
log10(ALDEx2Wilcoxon p-value)
log10(Wilcoxon p-value)
Significant by
Both (n=468)
Neither (n=28770)
Wilcoxon (n=7228)
(RNAseq-mouseDiet) Wilcoxon vs. ANCOMBC2
log10(ANCOMBC2 p-value)
log10(Wilcoxon p-value)
Significant by
ANCOMBC2 (n=1224)
Both (n=5356)
Neither (n=27546)
Wilcoxon (n=2340)
(RNAseq-mouseDiet) Wilcoxon vs. metagenomeSeq
log10(metagenomeSeq p-value)
log10(Wilcoxon p-value)
Significant by
Both (n=1438)
metagenomeSeq (n=800)
Neither (n=27970)
Wilcoxon (n=6258)
(RNAseq-mouseDiet) DESeq2 vs. ALDEx2t-test
log10(ALDEx2t-test p-value)
log10(DESeq2 p-value)
Significant by
ALDEx2t-test (n=3)
Both (n=425)
DESeq2 (n=5656)
Neither (n=30382)
(RNAseq-mouseDiet) DESeq2 vs. ALDEx2Wilcoxon
log10(ALDEx2Wilcoxon p-value)
log10(DESeq2 p-value)
Significant by
ALDEx2Wilcoxon (n=1)
Both (n=467)
DESeq2 (n=5614)
Neither (n=30384)
(RNAseq-mouseDiet) DESeq2 vs. ANCOMBC2
log10(ANCOMBC2 p-value)
log10(DESeq2 p-value)
Significant by
ANCOMBC2 (n=1703)
Both (n=4877)
DESeq2 (n=1204)
Neither (n=28682)
(RNAseq-mouseDiet) DESeq2 vs. metagenomeSeq
log10(metagenomeSeq p-value)
log10(DESeq2 p-value)
Significant by
Both (n=1235)
DESeq2 (n=4846)
metagenomeSeq (n=1003)
Neither (n=29382)
(RNAseq-mouseDiet) edgeR vs. ALDEx2t-test
log10(ALDEx2t-test p-value)
log10(edgeR p-value)
Significant by
ALDEx2t-test (n=1)
Both (n=427)
edgeR (n=5574)
Neither (n=30464)
(RNAseq-mouseDiet) edgeR vs. ALDEx2Wilcoxon
log10(ALDEx2Wilcoxon p-value)
log10(edgeR p-value)
Significant by
Both (n=468)
edgeR (n=5533)
Neither (n=30465)
(RNAseq-mouseDiet) edgeR vs. ANCOMBC2
log10(ANCOMBC2 p-value)
log10(edgeR p-value)
Significant by
ANCOMBC2 (n=1920)
Both (n=4660)
edgeR (n=1341)
Neither (n=28545)
(RNAseq-mouseDiet) edgeR vs. metagenomeSeq
log10(metagenomeSeq p-value)
log10(edgeR p-value)
Significant by
Both (n=1470)
edgeR (n=4531)
metagenomeSeq (n=768)
Neither (n=29697)

S17.

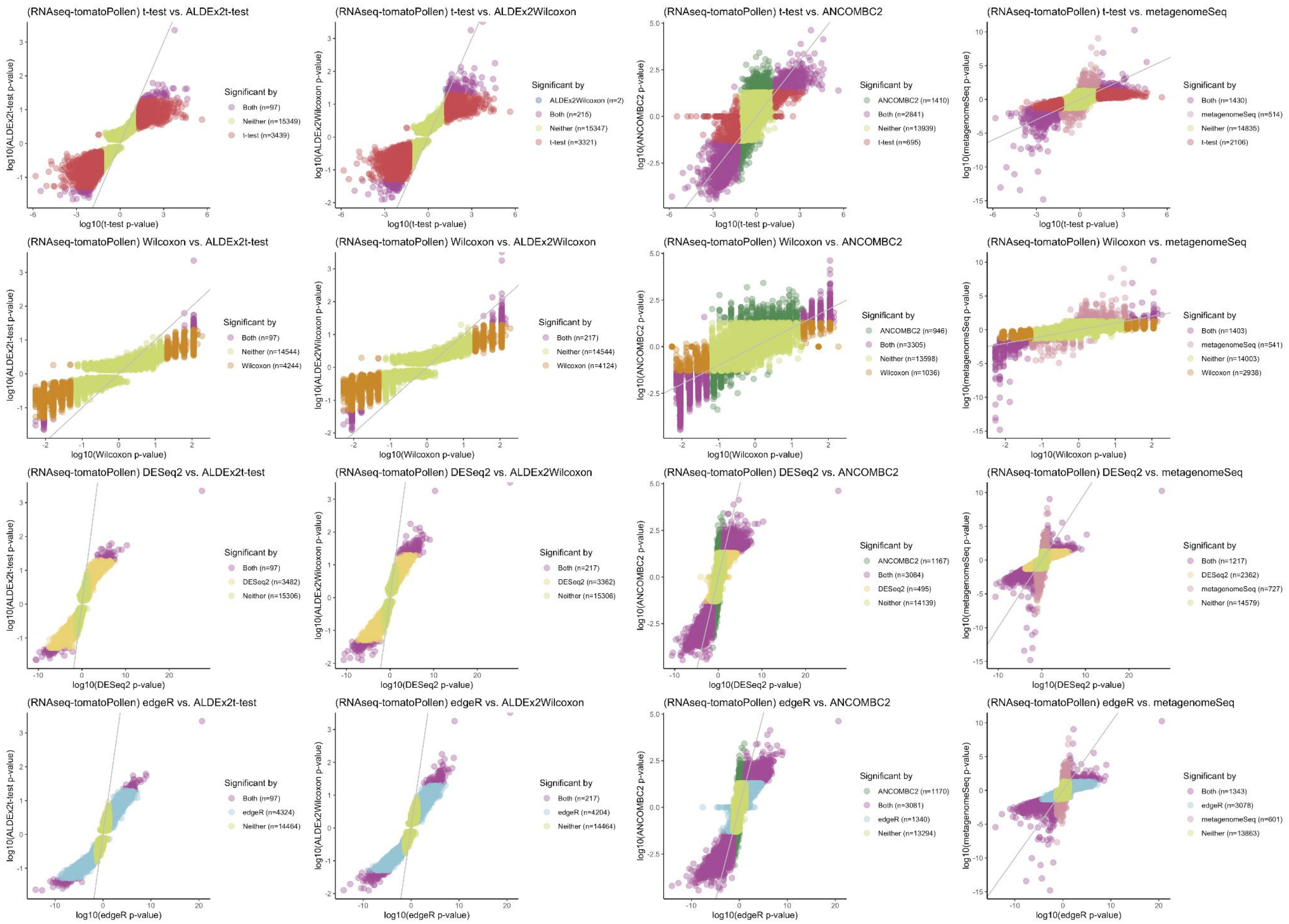
(RNAseq-tomatoPollen) t-test vs. ALDEx2t-test
log10(ALDEx2t-test p-value)
log10(t-test p-value)
Significant by
Both (n=97)
Neither (n=15349)
t-test (n=3439)
(RNAseq-tomatoPollen) t-test vs. ALDEx2Wilcoxon
log10(ALDEx2Wilcoxon p-value)
log10(t-test p-value)
Significant by
ALDEx2Wilcoxon (n=2)
Both (n=215)
Neither (n=15347)
t-test (n=3321)
(RNAseq-tomatoPollen) t-test vs. ANCOMBC2
log10(ANCOMBC2 p-value)
log10(t-test p-value)
Significant by
ANCOMBC2 (n=1410)
Both (n=2841)
Neither (n=13939)
t-test (n=695)
(RNAseq-tomatoPollen) t-test vs. metagenomeSeq
log10(metagenomeSeq p-value)
log10(t-test p-value)
Significant by
Both (n=1430)
metagenomeSeq (n=514)
Neither (n=14835)
t-test (n=2106)
(RNAseq-tomatoPollen) Wilcoxon vs. ALDEx2t-test
log10(ALDEx2t-test p-value)
log10(Wilcoxon p-value)
Significant by
Both (n=97)
Neither (n=14544)
Wilcoxon (n=4244)
(RNAseq-tomatoPollen) Wilcoxon vs. ALDEx2Wilcoxon
log10(ALDEx2Wilcoxon p-value)
log10(Wilcoxon p-value)
Significant by
Both (n=217)
Neither (n=14544)
Wilcoxon (n=4124)
(RNAseq-tomatoPollen) Wilcoxon vs. ANCOMBC2
log10(ANCOMBC2 p-value)
log10(Wilcoxon p-value)
Significant by
ANCOMBC2 (n=946)
Both (n=3305)
Neither (n=13598)
Wilcoxon (n=1036)
(RNAseq-tomatoPollen) Wilcoxon vs. metagenomeSeq
log10(metagenomeSeq p-value)
log10(Wilcoxon p-value)
Significant by
Both (n=1403)
metagenomeSeq (n=541)
Neither (n=14003)
Wilcoxon (n=2938)
(RNAseq-tomatoPollen) DESeq2 vs. ALDEx2t-test
log10(ALDEx2t-test p-value)
log10(DESeq2 p-value)
Significant by
Both (n=97)
DESeq2 (n=3482)
Neither (n=15306)
(RNAseq-tomatoPollen) DESeq2 vs. ALDEx2Wilcoxon
log10(ALDEx2Wilcoxon p-value)
log10(DESeq2 p-value)
Significant by
Both (n=217)
DESeq2 (n=3362)
Neither (n=15306)
(RNAseq-tomatoPollen) DESeq2 vs. ANCOMBC2
log10(ANCOMBC2 p-value)
log10(DESeq2 p-value)
Significant by
ANCOMBC2 (n=1167)
Both (n=3084)
DESeq2 (n=495)
Neither (n=14139)
(RNAseq-tomatoPollen) DESeq2 vs. metagenomeSeq
log10(metagenomeSeq p-value)
log10(DESeq2 p-value)
Significant by
Both (n=1217)
DESeq2 (n=2362)
metagenomeSeq (n=727)
Neither (n=14579)
(RNAseq-tomatoPollen) edgeR vs. ALDEx2t-test
log10(ALDEx2t-test p-value)
log10(edgeR p-value)
Significant by
Both (n=97)
edgeR (n=4324)
Neither (n=14464)
(RNAseq-tomatoPollen) edgeR vs. ALDEx2Wilcoxon
log10(ALDEx2Wilcoxon p-value)
log10(edgeR p-value)
Significant by
Both (n=217)
edgeR (n=4204)
Neither (n=14464)
(RNAseq-tomatoPollen) edgeR vs. ANCOMBC2
log10(ANCOMBC2 p-value)
log10(edgeR p-value)
Significant by
ANCOMBC2 (n=1170)
Both (n=3081)
edgeR (n=1340)
Neither (n=13294)
(RNAseq-tomatoPollen) edgeR vs. metagenomeSeq
log10(metagenomeSeq p-value)
log10(edgeR p-value)
Significant by
Both (n=1343)
edgeR (n=3078)
metagenomeSeq (n=601)
Neither (n=13863)

S18.

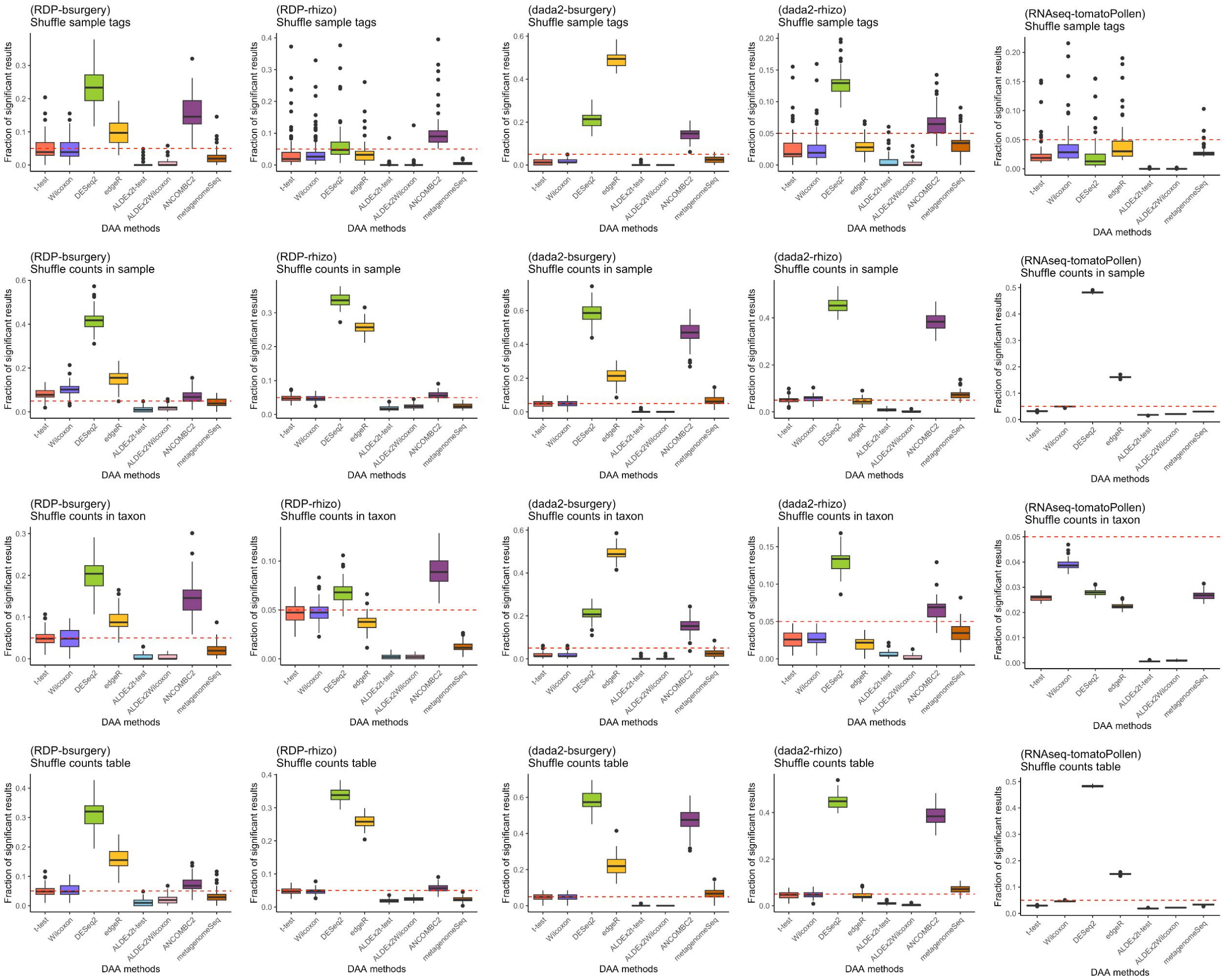
(RDP-bsurgery)
Shuffle sample tags
(RDP-rhizo)
Shuffle sample tags
(dada2-bsurgery)
Shuffle sample tags
(dada2-rhizo)
Shuffle sample tags
(RNAseq-tomatoPollen)
Shuffle sample tags
(RDP-bsurgery)
Shuffle counts in sample
(RDP-rhizo)
Shuffle counts in sample
(dada2-bsurgery)
Shuffle counts in sample
(dada2-rhizo)
Shuffle counts in sample
(RNAseq-tomatoPollen)
Shuffle counts in sample
(RDP-bsurgery)
Shuffle counts in taxon
(RDP-rhizo)
Shuffle counts in taxon
(dada2-bsurgery)
Shuffle counts in taxon
(dada2-rhizo)
Shuffle counts in taxon
(RNAseq-tomatoPollen)
Shuffle counts in taxon
(RDP-bsurgery)
Shuffle counts table
(RDP-rhizo)
Shuffle counts table
(dada2-bsurgery)
Shuffle counts table
(dada2-rhizo)
Shuffle counts table
(RNAseq-tomatoPollen)
Shuffle counts table
Fraction of significant results
DAA methods
t-test
Wilcoxon
DESeq2
edgeR
ALDEx2t-test
ALDEx2Wilcoxon
ANCOMBC2
metagenomeSeq

S19.

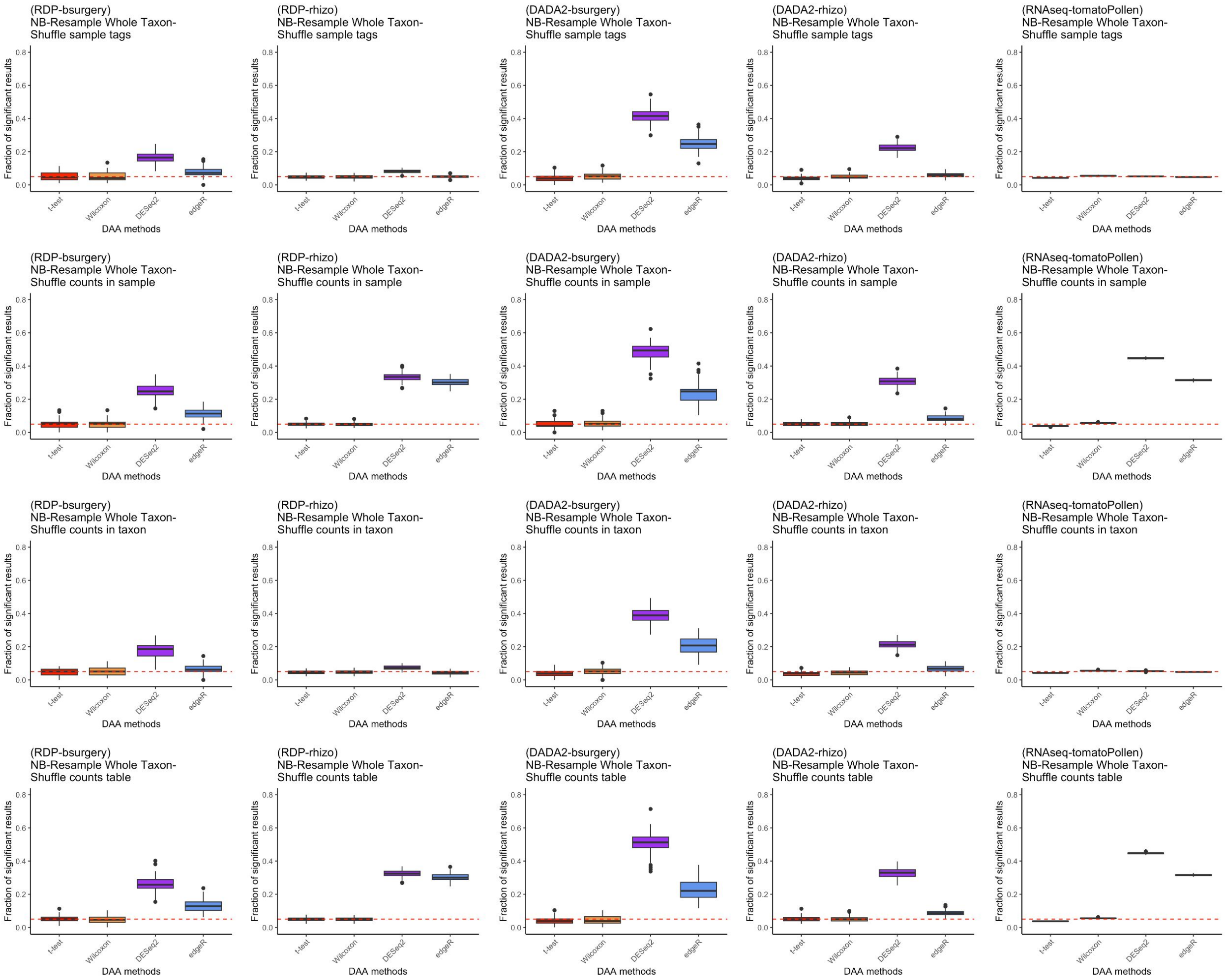
(RDP-bsurgery)
NB-Resample Whole Taxon-
Shuffle sample tags
(RDP-rhizo)
NB-Resample Whole Taxon-
Shuffle sample tags
(DADA2-bsurgery)
NB-Resample Whole Taxon-
Shuffle sample tags
(DADA2-rhizo)
NB-Resample Whole Taxon-
Shuffle sample tags
(RNAseq-tomatoPollen)
NB-Resample Whole Taxon-
Shuffle sample tags
(RDP-bsurgery)
NB-Resample Whole Taxon-
Shuffle counts in sample
(RDP-rhizo)
NB-Resample Whole Taxon-
Shuffle counts in sample
(DADA2-bsurgery)
NB-Resample Whole Taxon-
Shuffle counts in sample
(DADA2-rhizo)
NB-Resample Whole Taxon-
Shuffle counts in sample
(RNAseq-tomatoPollen)
NB-Resample Whole Taxon-
Shuffle counts in sample
(RDP-bsurgery)
NB-Resample Whole Taxon-
Shuffle counts in taxon
(RDP-rhizo)
NB-Resample Whole Taxon-
Shuffle counts in taxon
(DADA2-bsurgery)
NB-Resample Whole Taxon-
Shuffle counts in taxon
(DADA2-rhizo)
NB-Resample Whole Taxon-
Shuffle counts in taxon
(RNAseq-tomatoPollen)
NB-Resample Whole Taxon-
Shuffle counts in taxon
(RDP-bsurgery)
NB-Resample Whole Taxon-
Shuffle counts table
(RDP-rhizo)
NB-Resample Whole Taxon-
Shuffle counts table
(DADA2-bsurgery)
NB-Resample Whole Taxon-
Shuffle counts table
(DADA2-rhizo)
NB-Resample Whole Taxon-
Shuffle counts table
(RNAseq-tomatoPollen)
NB-Resample Whole Taxon-
Shuffle counts table
Fraction of significant results
0.0
0.2
0.4
0.6
0.8
t-test
Wilcoxon
DESeq2
edgeR
DAA methods

S20.

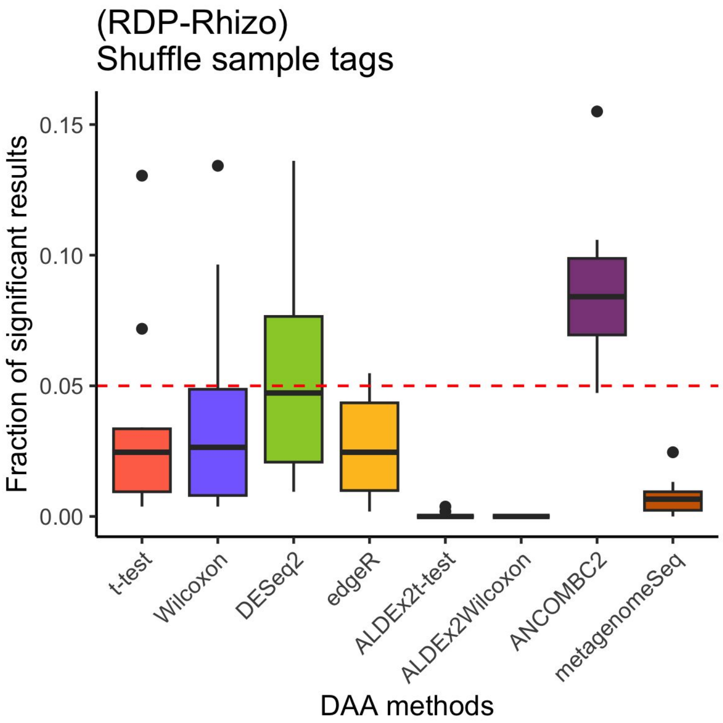

(RDP-Rhizo)
Shuffle sample tags
Fraction of significant results
0.15
0.10
0.05
0.00
t-test
Wilcoxon
DESeq2
edgeR
ALDEx2t-test
ALDEx2Wilcoxon
ANCOMBC2
metagenomeSeq
DAA methods


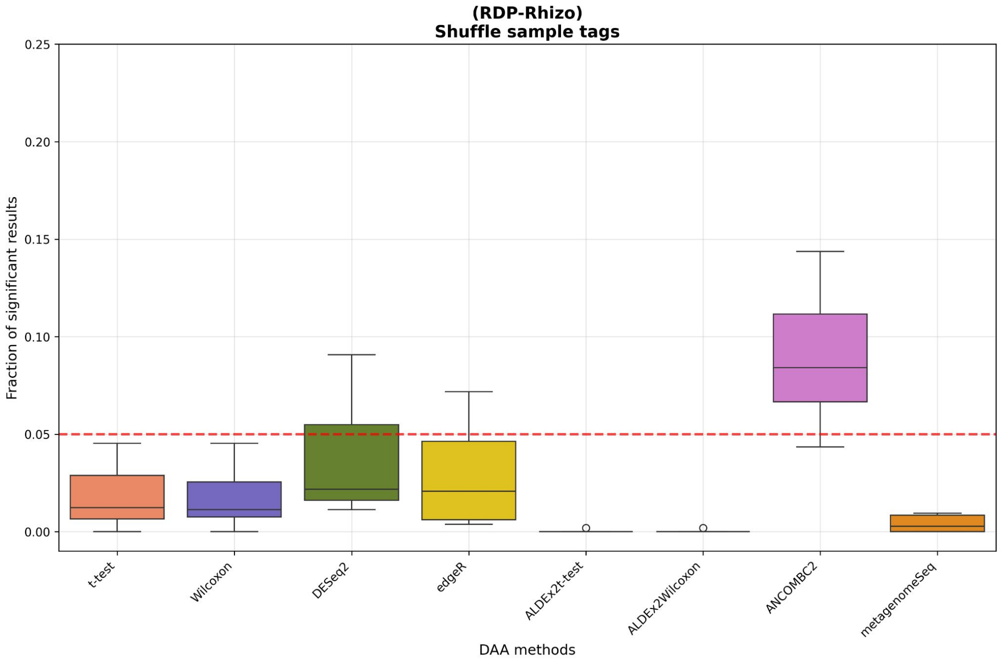

(RDP-Rhizo)
Shuffle sample tags
Fraction of significant results
0.25
0.20
0.15
0.10
0.05
0.00
t-test
Wilcoxon
DESeq2
edgeR
ALDEx2t-test
ALDEx2Wilcoxon
ANCOMBC2
metagenomeSeq
DAA methods
